# Transcriptome-based phylogeny of the semi-aquatic bugs (Hemiptera: Heteroptera: Gerromorpha) reveals patterns of lineage expansion in a series of new adaptive zones

**DOI:** 10.1101/2022.01.08.475494

**Authors:** David Armisén, Séverine Viala, Isabelle da Rocha Silva Cordeiro, Antonin Jean Johan Crumière, Elisa Hendaoui, Augustin Le Bouquin, Wandrille Duchemin, Emilia Santos, William Toubiana, Aidamalia Vargas-Lowman, Carla Fernanda Burguez Floriano, Dan A. Polhemus, Yan-hui Wang, Locke Rowe, Felipe Ferraz Figueiredo Moreira, Abderrahman Khila

## Abstract

Key innovations enable access to new adaptive zones and are often linked to increased species diversification. As such, they have attracted much attention, yet their concrete consequences on the subsequent evolutionary trajectory and diversification of the bearing lineages remain unclear. The monophyletic group of water striders and relatives (Hemiptera: Heteroptera: Gerromorpha) represent a group of insects that transited to live on the water-air interface and diversified to occupy ponds, puddles, streams, mangroves and even oceans. This lineage offers an excellent model to study the patterns and processes underlying species diversification following the conquest of new adaptive zones. However, such studies require a reliable and comprehensive phylogeny of the infraorder. Based on whole transcriptomic datasets of 97 species and fossil records, we reconstructed a new phylogeny of the Gerromorpha that resolved inconsistencies and uncovered strong support for previously unknown relationships between some important taxa. We then used this phylogeny to reconstruct the ancestral state of a set of adaptations associated with water surface invasion (fluid locomotion, dispersal and transition to saline waters) and sexual dimorphism. Our results uncovered important patterns and dynamics of phenotypic evolution revealing how the initial event of water surface invasion enabled multiple subsequent transitions to new adaptive zones, representing distinct niches of water surfaces, and further diversification of the group. This phylogeny and the associated transcriptomic datasets constitute highly valuable resources, making Gerromorpha an attractive model lineage to study phenotypic evolution.

## Introduction

Key innovations enable access to new adaptive zones where lineages can diversify and occupy new niches (Simpson 1953; Mayr 1963; Schluter 2000; Rabosky 2017). These innovations have attracted much attention, because they are often linked to increased diversification (Liem 1973; Schluter 2000; Wagner and Lynch 2010; Wagner 2017; Ronco, et al. 2021). Spectacular examples of this process are thought to include the evolution of flight in bats and herbivory in insects. Understanding the processes underlying transitions to new adaptive zones and the consequences of these transitions on the evolutionary trajectory of lineages has greatly enriched our understanding of species diversification. These studies typically focus on the evolution and ecological function of traits or sets of traits that constitute the innovation and its consequences for diversity in the lineage. More recently, studies have begun to focus on the developmental genetics underlying these traits (Abzhanov, et al. 2004; Santos, et al. 2017).

Arthropods inhabit a great span of habitats, from marine to freshwater, and from land to air. However, very few lineages permanently inhabit the air-water interface (e.g., whirligig beetles, fishing spiders and water striders). This life style may be limited by the constraints of fluid dynamics on remaining on the water surface and generating efficient thrust for movement on the fluid substrate (Andersen 1982; Hu and Bush 2010). Here we focus on water striders and relatives, also known as semi-aquatic bugs (Hemiptera: Heteroptera: Gerromorpha), which represent a group of insects characterized by their ability to live on the air-water interface (Andersen 1982). This group is a prominent example of a lineage that transited to a new adaptive zone, the water surface, and offers a unique opportunity to study the causes and consequences of this transition on species diversification. Since the initial transition to water surface habitats, the semi-aquatic bugs radiated into over 2,000 extant species occupying a vast array of niches ranging from rain puddles to ponds, streams, lakes, mangroves, and even the open oceans (Andersen 1982; Polhemus and Polhemus 2008).

The gerromorphans are characterized by a large array of adaptive, and often conspicuous, traits associated with water surface locomotion. A key evolutionary event that characterizes the history of this lineage is the original invasion of water surface habitats from a common terrestrial ancestor dating to over 200 MYA (Damgaard 2008b; Wang, et al. 2016; Johnson, et al. 2018). The transition to live on the water is linked to the ability of water striders to support their body weight on the water surface by exploiting surface tension (Andersen 1976; Hu and Bush 2010). Layers of hydrophobic hairs covering their legs sequester air bubbles, forming a cushion between the legs and the water, thus allowing the animals to avoid breaking the surface tension (Andersen 1982; Gao and Jiang 2004; Hu and Bush 2010). Other traits represent adaptations to generating efficient thrust on the fluid substrate. These include the elongation of the legs relative to terrestrial species that increases speed on the water (Crumière, et al. 2016). This change is characteristic of lineages that occupy marginal aquatic zones and that can walk both on water and on land (Andersen 1982; Crumière, et al. 2016). Other lineages have specialized in open water zones and propel their body by means of surface rowing (Andersen 1982). This derived mode of locomotion is tightly associated with a change in the body plan to a derived state where midlegs are longer and are primarily responsible for propulsion through simultaneous sculling motion, also known as the rowing gait (Andersen 1976). While the contribution of the set of adaptive traits to the initial transition to the water surface has been subject of many studies (Andersen 1982; Gao and Jiang 2004; Hu and Bush 2010; Wang, et al. 2015), the consequences of this transition on the subsequent evolutionary trajectories and diversification of the lineage remain unclear.

Gerromorpha also represents a prominent model for the study of sexual selection and sexual conflict, *e.g.* (Rowe, et al. 1994; Fairbairn and Preziosi 1996; Arnqvist 1997; Preziosi and Fairbairn 2000; Arnqvist and Rowe 2002a, b; Ronkainen, et al. 2010; Crumière and Khila 2019; Toubiana and Khila 2019; Toubiana, Armisén, Dechaud, et al. 2021). In many species, the mating system is characterized by sexual conflict, which is often manifested by rigorous pre-mating struggles where females resist costly mating attempts and males persist (Arnqvist 1989; Rowe 1994; Ronkainen, et al. 2010). These premating struggles have favored the repeated evolution of remarkable grasping structures in males and anti-grasping structures in females (Rowe, et al. 2006; Crumière, et al. 2019), which appear to coevolve antagonistically (Arnqvist and Rowe 2002a; Perry, et al. 2017; Crumière and Khila 2019). It is interesting that the characteristic sexual conflict in the mating system of many members of this group may itself have resulted from the transition to the 2-dimensional water surface habitat, where females are relatively easy to detect, locate and approach (Arnqvist 1997).

Previous works documented the developmental genetic mechanisms underlying appendage diversification in a selection of gerromorphans. The Hox gene *Ultrabithorax* has been key to increasing leg length and in changes in body plan associated with water surface locomotion and predation escape (Khila, et al. 2009, 2014; Refki, et al. 2014; Armisén, et al. 2015; Refki and Khila 2015). Interestingly, the same gene is involved in modifying male third legs into claspers under sexually antagonistic selection in the water strider *Rhagovelia antilleana* (Crumière and Khila 2019). In addition, we recently began to elucidate the developmental genetics of other sexually antagonistic traits consisting of modified antennae (Khila, et al. 2012) or other independent cases of male third leg modifications (Toubiana, Armisén, Dechaud, et al. 2021; Toubiana, Armisén, Viala, et al. 2021). However, the evolutionary history and the combined impact of water surface locomotion and sexual conflict on the diversification of the legs in water striders remain to be tested.

This rich natural history, phenotypic diversity, and the development of genetic tools, together with the invasion of the water surface habitat, provide an invaluable opportunity to study the patterns and processes underlying phenotypic evolution of lineages following the transition to new adaptive zones. A major constraint on this opportunity is the lack of a well supported phylogeny of the whole infraorder, since those currently available have been recovered using a limited number of morphological characters and/or molecular markers (*e.g.* (Andersen 1982; Andersen and Weir 2004; Damgaard 2008a). While the current phylogenies generally agree on the position of certain nodes at higher taxonomic levels, some lineages remain problematic or unresolved (Damgaard 2008a). In particular, the currently accepted higher classification of the group has inconsistencies at the levels of family and subfamily, and key taxa remain unassessed (Damgaard 2008a; Damgaard 2012; Damgaard, et al. 2014; Román-Palacios, et al. 2020). These inconsistencies hinder reliable reconstructions of ancestral character states, thus limiting our understanding of the evolutionary history of this group.

In this work, we sampled, sequenced and *de novo* assembled the full transcriptomes of ninety-seven species of Gerromorpha. We identified a set of 894 common molecular markers that we used to reconstruct the phylogeny of the infraorder. We resolved inconsistencies and uncovered strong support for previously unknown relationships between some important taxa. We then calibrated the phylogeny based on the available fossil record. Finally, we used this phylogeny to reconstruct the evolutionary history of a set of adaptive characters associated with water surface locomotion, dispersal, transition from fresh to saline waters and sexual dimorphism. Our results reveal how the initial event of water surface invasion enabled multiple subsequent transitions to new adaptive zones and further diversification of the group.

## Results

### Transcriptome assemblies and gene clustering

We sequenced the transcriptomes of 95 species, of which 92 cover five out of the eight known families of Gerromorpha and 3 are outgroup species (Supplementary Material Table 1). The number of Illumina reads obtained ranged between 43,134,196 and 274,421,352 (Supplementary Material Table 2). Transcriptome assemblies using Trinity yielded between 52,462 and 363,535 transcripts with BUSCOs 5.2.2 completeness ranging from 30.7 % to 95.1 % (Figure 1 and Supplementary Material Table 2). Nine additional transcriptomes from other Gerromorpha or closely related species were included from other sources (see material and methods). A total of 104 species (97 Gerromorpha + 7 outgroup species) were used to reconstruct the phylogeny. Clustering of the combined 3,299,629 transcriptomic genes in all species yielded a total of 3,079,991 clusters, including singletons. Despite the high initial number of clusters, due to our stringency criteria and the high number of species, this step reduced the list to 3,169 clusters of transcripts that are common in at least 80% of the species to reconstruct the phylogeny of Gerromorpha (see methods). After testing for stationary, reversible, and homogeneous (SRH) consistency, a final list of 894 gene clusters was retained for phylogeny reconstruction using IQTREE (see methods) (Supplementary Figure 1).

**Figure 1:**
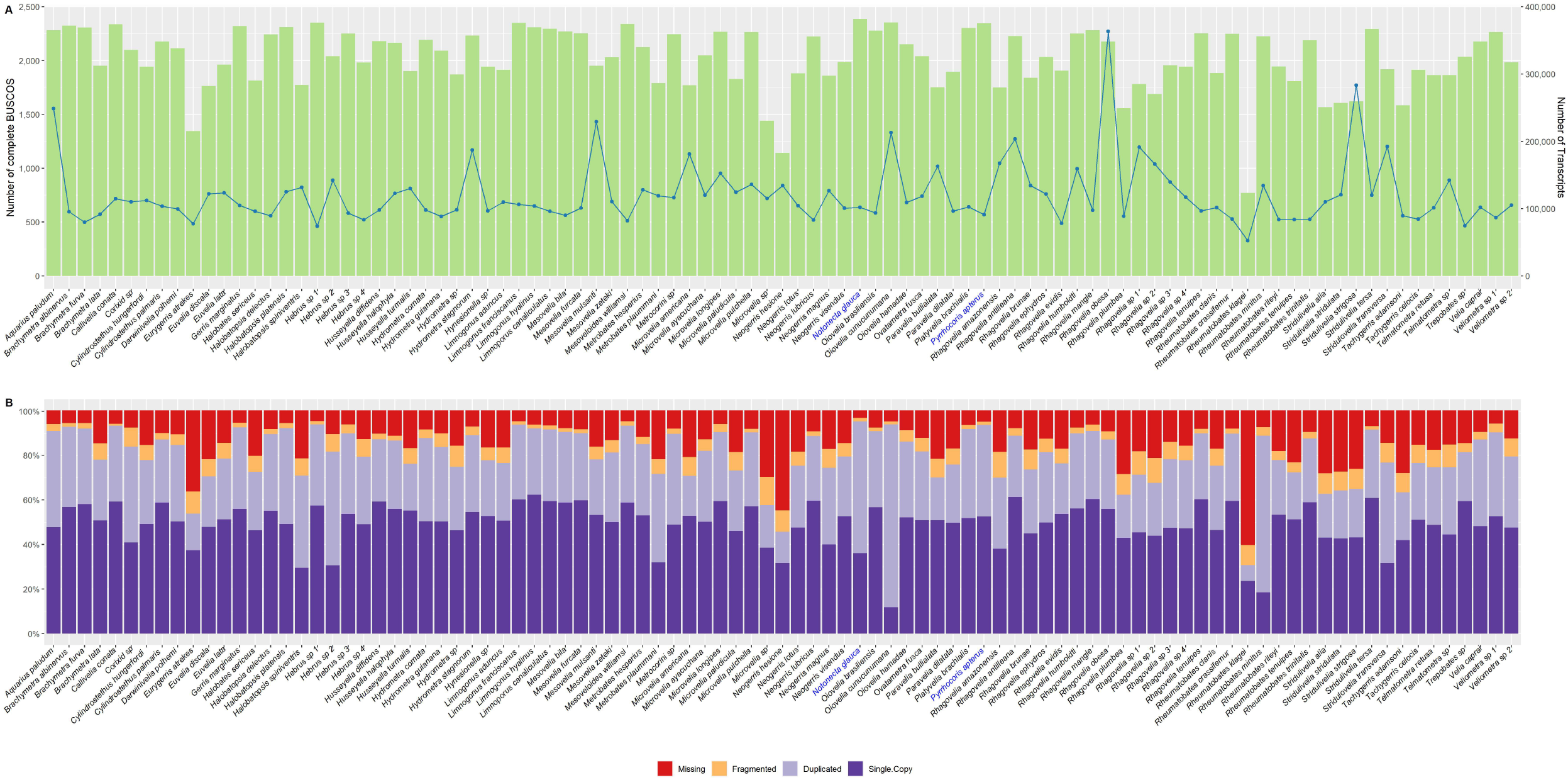
(A) Green histogram: Number of complete BUSCOS found in the assembled transcriptome of each newly sequenced species (92 Gerromorpha and 3 outgroup species); Blue line: Number of *de novo* assembled transcripts by Trinity. (B) Breakdown of BUSCOS results percentages including complete (single-copy in dark purple and duplicated in light purple), fragmented in orange and missing in red. In both cases names of outgroup species (i.e. not Gerromorpha) were colored in blue.

### Recovering Gerromorpha phylogeny using transcriptomic markers

The infraorder Gerromorpha, as well as the families Mesoveliidae, Hebridae, and Gerridae, were all recovered as monophyletic (Figure 2). Hydrometridae (hereafter “Hydrometridae”) is however polyphyletic, because *Veliometra* (“Hydrometridae”: Heterocleptinae) is closer to *Hebrus* (Hebridae: Hebrinae) than to *Hydrometra* (“Hydrometridae”: Hydrometrinae). Because this is the first time that this relationship is recovered and we lack samples of other “Hydrometridae” genera except *Hydrometra*, we refrain from proposing any classification changes at this moment. Mesoveliidae is sister to (Hebridae + *Veliometra*), the three are sister to *Hydrometra*, and the clade as a whole is sister to (Veliidae + Gerridae). Veliidae (from now on “Veliidae”) is also not monophyletic (paraphyletic), consisting of a succession of clades sister to Gerridae. The same is also true for the subfamily Veliinae (“Veliidae”), in which *Velia caprai* was recovered as sister to all other clades of (“Veliidae” + Gerridae), while a clade containing species of *Callivelia, Oiovelia,* and *Paravelia* is sister to *Rhagovelia* (“Veliidae”: Rhagoveliinae), not to the rest of the subfamily. Furthermore, the clade (Microveliinae + Haloveliinae) is more closely related to Gerridae than to the other subfamilies of “Veliidae”. Within Gerridae in the current sense (e.g. Damgaard 2008), Halobatinae was recovered as sister to all other subfamilies, and Rhagadotarsinae as sister to (Trepobatinae + (Gerrinae + (Charmatometrinae + Cylindrostethinae))).

**Figure 2:**
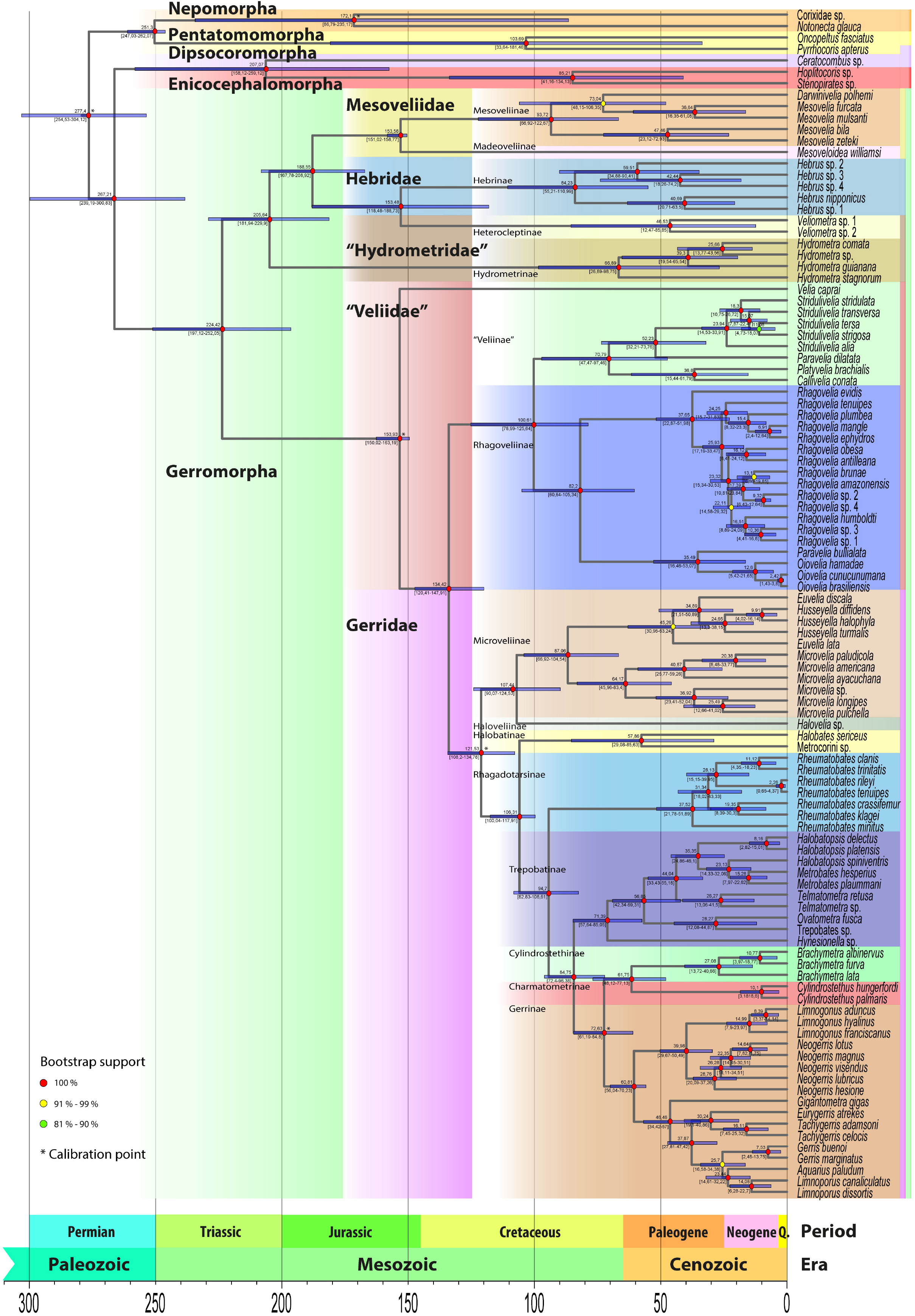
Dated phylogeny of Gerromorpha. Timescale was estimated from 5 calibration points (marked with an asterisk (*)). Estimated ages are indicated on each node and the 95% confidence interval is indicated with a blue box while confidence interval ages are presented in between square brackets. Families are highlighted in various colors with family names indicated.

Among the results above, the sister relationship between Gerridae and (Microveliinae + Haloveliinae) had already been detected by Damgaard (Damgaard 2008a), who did not propose any classification changes. The whole clade was recovered here with full support and has two strong morphological synapomorphies. The first is the fusion of tarsomeres I and II in all legs that results in the tarsal formula 2:2:2 in Haloveliinae and Gerridae, which is further reduced to 1:2:2 by the fusion of all foreleg tarsomeres in Microveliinae (Andersen 1982). The second is the presence of a fecundation pump on the female gynatrial complex, which is believed to assist in the ejection of sperm through the apical part of the fecundation canal and into the lumen of the common oviduct prior to the fertilization of the passing egg (Andersen 1982). These observations indicate that Microveliinae and Haloveliinae are not veliids, which leads us to propose transferring them to the family Gerridae. Even with the removal of these two subfamilies, the current classification of “Veliidae” remains highly artificial and in need of a thorough revision.

### Time of divergence of the main clades of Gerromorpha

To date the divergence time in the phylogenetic tree of Gerromorpha, we used five fossil calibration points obtained from eight fossil records (Supplementary Material Table 3). The split of Gerromorpha was estimated to have occurred during the mid Permian about 267 million year ago (MYA) (95% CI 239.19-300.83 Mya) (Figure 2), consistent with other phylogenetic analyses (Wang, et al. 2016; Johnson, et al. 2018; Wang, et al. 2021). The divergence of the major clades of Gerromorpha is estimated to have started during the mid Triassic about 224.42 MYA (95% CI 197.12-252.05 Mya) (Figure 2) when the clade (Gerridae + “Veliidae”) split from (“Hydrometridae” + (Mesoveliidae + Hebridae)) early during the evolution of Gerromorpha.

This result is inconsistent with classic phylogenies using few markers that recovered (Gerridae + Veliidae) as a derived and later branching clade (Andersen and Weir 2004; Damgaard 2008a), but consistent with most recent phylogenetic analyses using larger datasets (Wang, et al. 2016; Johnson, et al. 2018; Wang, et al. 2021). In this scenario, “Hydrometridae” forms a sister clade to (Hebridae + Mesoveliidae), from which it split 205 MYA (95% CI 181.94-229.9 Mya), whereas the Hebridae split from Mesoveliidae about 188 MYA (95% CI 167.78-208.92 Mya). Also, contrary to previous reconstructions clustering *Veliometra* within the Hydrometridae based on morphology alone (Andersen 1982; Andersen and Weir 2004), *Veliometra* is sister to *Hebrus* and the two clades split about 153 MYA (95% CI 118.48-188.73 Mya).

On the other side of the tree, the split between “Veliidae” (already excluding Microveliinae and Haloveliinae as explained in the previous section) and Gerridae took place during the Mesozoic, about 153 MYA (95% CI 150.02-163.19 Mya) (Figure 2). In turn, Microveliinae + Haloveliinae diverged from the other Gerridae subfamilies about 134 MYA (95% CI 120.41-147.91 Mya).

### Single origin of rowing as a derived mode of locomotion in “Veliidae” and Gerridae

Ancestral Gerromorpha transited from terrestrial to a new adaptive zone at the margins of the water in a single event. Subsequent to this transition, species occupy a vast and heterogeneous array of water surface niches (Andersen 1982). Lineages occupying the margins, the ancestral adaptive zone, employ the walking gait, with contribution from all three legs, and retain the ancestral body plan with hindleg being the longest (Andersen 1976; Crumière, et al. 2016). We have previously shown a strong and positive correlation between the length of midlegs and locomotion speed on water (Crumière, et al. 2016). Ancestral character state reconstruction revealed that, at the base of Gerromorpha and coinciding with the transition to water surface habitats, there was a significant increase in the ratio midleg to body length in both males and females (Supplementary Figures 3 and 6). The increase in leg length is more modest for the forelegs and hindlegs (Supplementary Figures 2, 4, 5 and 7), consistent with the primary role of the midlegs in generating propulsion even when the animals retain the ancestral walking gait (Crumière, et al. 2016). This data suggest that increased leg length is among the phenotypic changes associated with the acquisition of marginal water areas as a new adaptive zone.

Most lineages of Gerromorpha transited to another adaptive zone consisting of a variety of open waters, ranging from ponds and stream to oceans (Andersen 1982). These lineages are characterized by a derived mode of locomotion based on the rowing gait, which almost exclusively relies on the midlegs to generate thrust (Andersen 1976; Hu, et al. 2003; Crumière, et al. 2016). A biomechanical analysis established a strict connection between body plan and locomotion mode, whereby species with midlegs being the longest use rowing and those with hindlegs being the longest use the walking gait (Andersen 1976; Khila, et al. 2014; Crumière, et al. 2016). We therefore scored the state of body plan across our sample to reconstruct the evolutionary history of locomotion modes in Gerromorpha (Figure 3). We found that leg morphology associated with the walking gait is ancestral and that the switch to midlegs being longer evolved only once some 153 MYA in the common ancestor of (“Veliidae” + Gerridae) (Figure 3). Within this clade, we detected four independent reversals from the rowing morphology back to the walking morphology, three times in the “Veliidae’ and once in the “Microveliinae” (Figure 3). Interestingly, all these reversals in morphology were accompanied by reversals of habitat preference back to the margins of water bodies (Andersen 1976, 1982). The genus *Oiovelia* further specialized in foam, which accumulates at the margins of rivers or is trapped by fallen tree trunks or other obstacles, where they build tunnels for breeding (Rodrigues, et al. 2014). Combined with the evolutionary history of Gerromorpha, these results suggest that the rowing mode of locomotion represents an early adaptation that evolved shortly after the transition to life on the water surface. This early adaptation opened yet another adaptive zone consisting of the vast and diverse surfaces of open waters, and is associated with the impressive expansion of the clade (“Veliidae” + Gerridae), which accounts for about 80% of all gerromorphans (Polhemus and Polhemus 2008).

**Figure 3:**
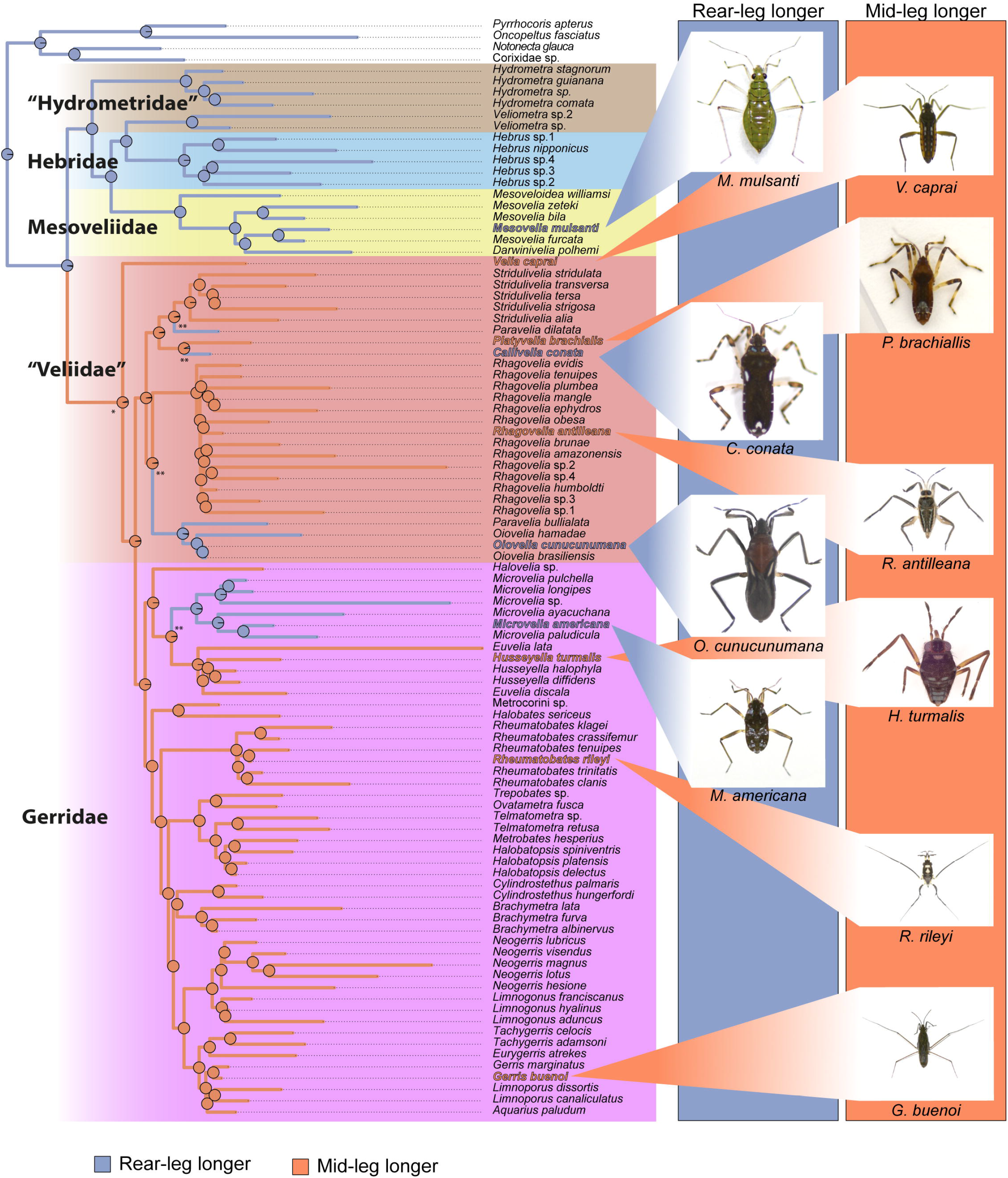
Ancestral character state reconstruction of relative leg length, with midlegs being the longest associated with rowing and hindlegs being the longest associated with walking (See (Crumière, et al. 2016). Pies represent the probability of ancestral state. Images represent samples of the two relative leg length phenotypes and associated locomotion mode. Derived body plan event and transition to open waters is marked with an asterisk (*). Reversals to ancestral body plan are marked with a double asterisk (**).

### Repeated co-option of male hindlegs into grasping structures through common genetic pathway

The biomechanical requirements of movement on the water surface appear to have driven a great deal of evolution of relative leg lengths in gerromorphans (Andersen 1982; Khila, et al. 2014; Crumière, et al. 2016). Sexually antagonistic coevolution has also resulted in elaboration and dimorphism of these appendages, sometimes profound (Rowe, et al. 2006; Crumière, et al. 2019). Some of the most common grasping traits in males are the modifications of the appendages into clamps that increase male’s ability to overcome female resistance to mating. Modification of the mid- and forelegs to increase their utility as clamps are common in the veliids and gerrids, yet modification of the midlegs into claspers, the primary appendage for water surface locomotion, is rare. Phylogenetic reconstruction revealed that the elaboration of males’ hindlegs into grasping traits evolved at least seven times independently in our sample (Figure 4). In some genera such as *Rheumatobates* or *Microvelia*, male modifications of the hindlegs evolved multiple times within the genus and are represented twice in our sampling (Figure 4). This is an under-estimation, as Rowe and colleagues have shown that male modified hindlegs evolved four times independently within the genus *Rheumatobates* (Rowe, et al. 2006). The modifications resulted in either divergent (*Rheumatobates*) or convergent (e. g. *Microvelia, Rhagovelia, Stridulivelia*) morphologies. Among the seven lineages that independently evolved modifications of male hindlegs, the females of five do not bear any modifications, suggesting frequent resolution of intra-locus sexual conflict (Stewart, et al. 2010). In the two genera *Stridulivelia* and *Rhagovelia* (except *R. plumbea* in our sample), females retain hindleg modifications albeit to a lesser degree than males, suggesting that genetic correlation between the sexes may constrain the evolution of sexual dimorphism in these two taxa (Figure 4). Interestingly, modified male hindlegs evolved only after the evolutionary elongation of the midlegs associated with rowing – a mode of locomotion that relies primarily on the midlegs (Andersen 1976; Hu et al. 2003; Crumière et al. 2016;). These results suggest that the primary use in locomotion of the midlegs may constrain their elaboration, whereas hindlegs (and forelegs) are freer to evolve in response to sexual conflict.

**Figure 4:**
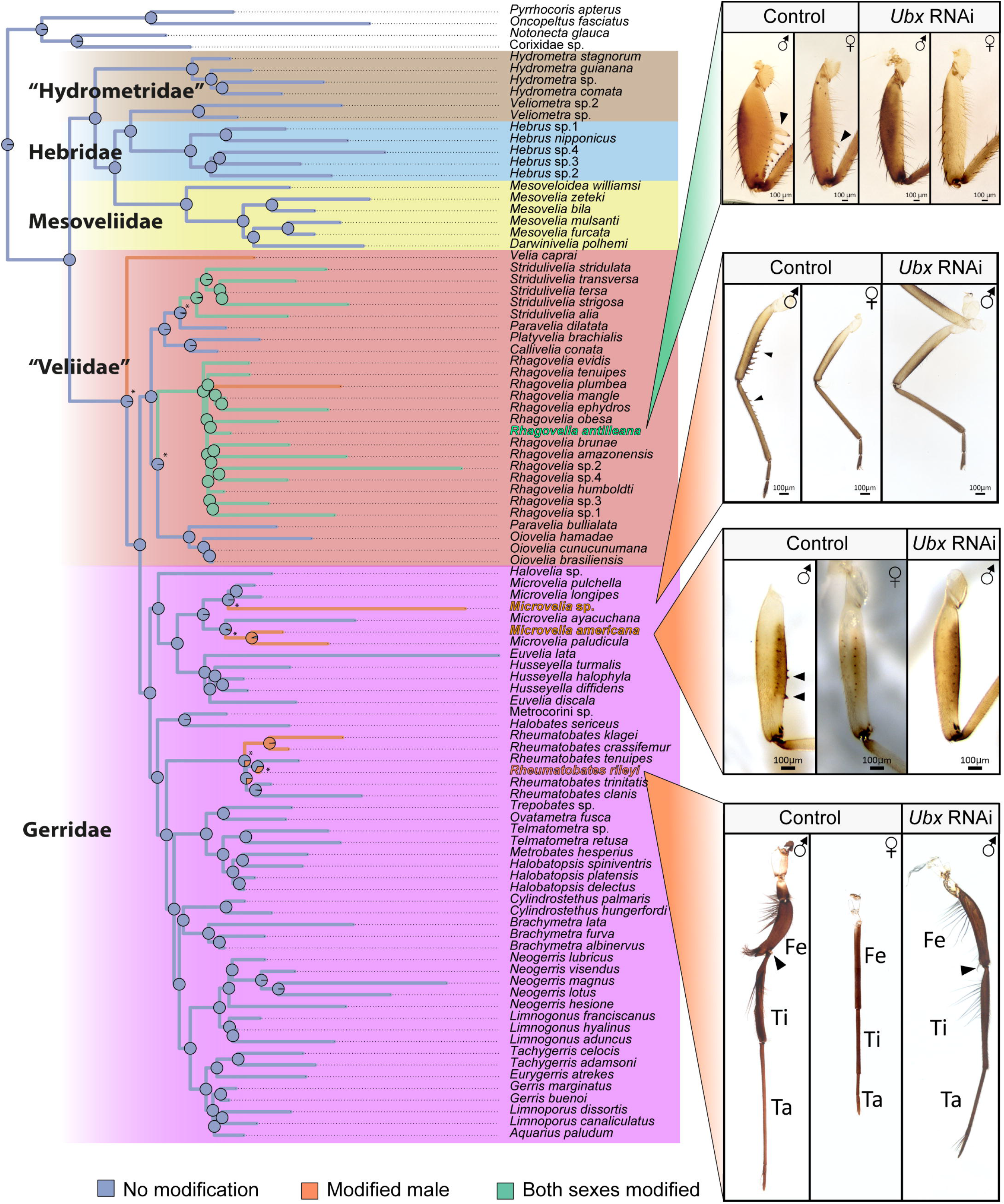
Ancestral character state reconstruction of male hindleg modifications into grasping traits driven by sexual conflict. Pies represent the probability of ancestral state. Images of four lineages that evolved male hindleg modifications, along with female hindleg comparisons, are represented. *Ubx* RNAi removes male modification in all lineages regardless of morphology or phylogenetic position. Independent leg modification events are marked with an asterisk (*).

We have previously shown that the Hox gene *Ultrabithorax* (*Ubx*) has a key role in leg shapes and relative sizes in Gerromorpha. Ubx is involved in the increase of the absolute length of both mid- and hindlegs in the walking species (Khila, et al. 2014; Refki, et al. 2014), in reversing the relative lengths of the mid- and hindlegs (and their segments) in the rowing species (Khila, et al. 2009, 2014; Refki, et al. 2014; Armisén, et al. 2015), and for the modifications of male hindlegs into clamps in the water strider *Rhagovelia antilleana* (Crumière and Khila 2019). Furthermore, *Ubx* depletion removes the modifications from females’ hindlegs, thus confirming that *R. antilleana* males and females share the genetic basis of hindleg modifications (Crumière and Khila 2019) (Figure 4). To test whether Ubx was involved in the other independent events of male hindleg modifications, we depleted *Ubx* transcripts using RNAi in three additional species, each representing an independent event of hindleg modifications in males (*Microvelia americana*, *Microvelia* sp. (new) and *Rheumatobates rileyi*). *Rheumatobates riley* males present hindleg modifications that are deeply divergent from *Rhagovelia antilleana* and *Microvelia* spp. (Figure 4). Strikingly, *Ubx* depletion across these three additional species resulted in the loss of the modifications of male’s hindlegs regardless of the morphology of the leg or the phylogenetic position of the bearing species (Figure 4). These findings support the repeated and independent co-option of *Ubx* during the evolution of male sexually antagonistic traits driven by sexual conflict throughout the evolutionary history of the semi-aquatic bugs.

### Independent transitions from fresh to saline waters

Out of the million or so described species of insects, Gerromorpha have been astonishingly successful in colonizing saline water habitats, including the open oceans (Andersen 1982; Cheng 1989; Ryland and Tyler 1989; Andersen 1999; Ikawa, et al. 2012). Various lineages occupy waters with different degrees of salinity ranging from brackish to high salinity seawaters. The salty *versus* freshwater preference covers both micro- and macro-evolutionary scales (Andersen 1999). In some genera, both salty and freshwater populations of the same species can be found, but some lineages either at the family level or at the genus level are exclusively marine (Andersen 1999; Ikawa, et al. 2012). Our sampling included 11 species, out of 97, which were know to inhabit and were collected in marine or brackish environments (Figure 5). The ability to occupy salty water bodies evolved early in the Gerromorpha during the Cretaceous in the Mesoveliidae, “Veliidae”, and Gerridae (Figures 2 and 5). Our character state reconstruction revealed that, of these 10 lineages, seven have independently transited from fresh to saline environments (Figure 5). Some of these transitions occurred below the genus level, as is the case for *Rheumatobates, Rhagovelia* or *Mesovelia* (Figure 5). Other events characterize entire lineages; such is the case for *Halobates* or *Halovelia*. Among these transitions, some lineages specialize in mangroves where water salinity fluctuates from almost fresh to entirely marine within the same day. These include *Rheumatobates trinitatis, Husseyella halophyla, Rhagovelia mangle* and *Rhagovelia ephydros*. This indicates that these lineages are unspecialized and have evolved phenotypic plasticity that allows them to expand their habitat range to environments with short-term fluctuations in salinity. Other lineages, including *Halobates* or *Halovelia*, are exclusively marine, except for secondary reversions to the freshwater life (Román-Palacios, et al. 2020), indicating that they have become specialized on salt water (Cheng 1976; Sekimoto, et al. 2014).

**Figure 5:**
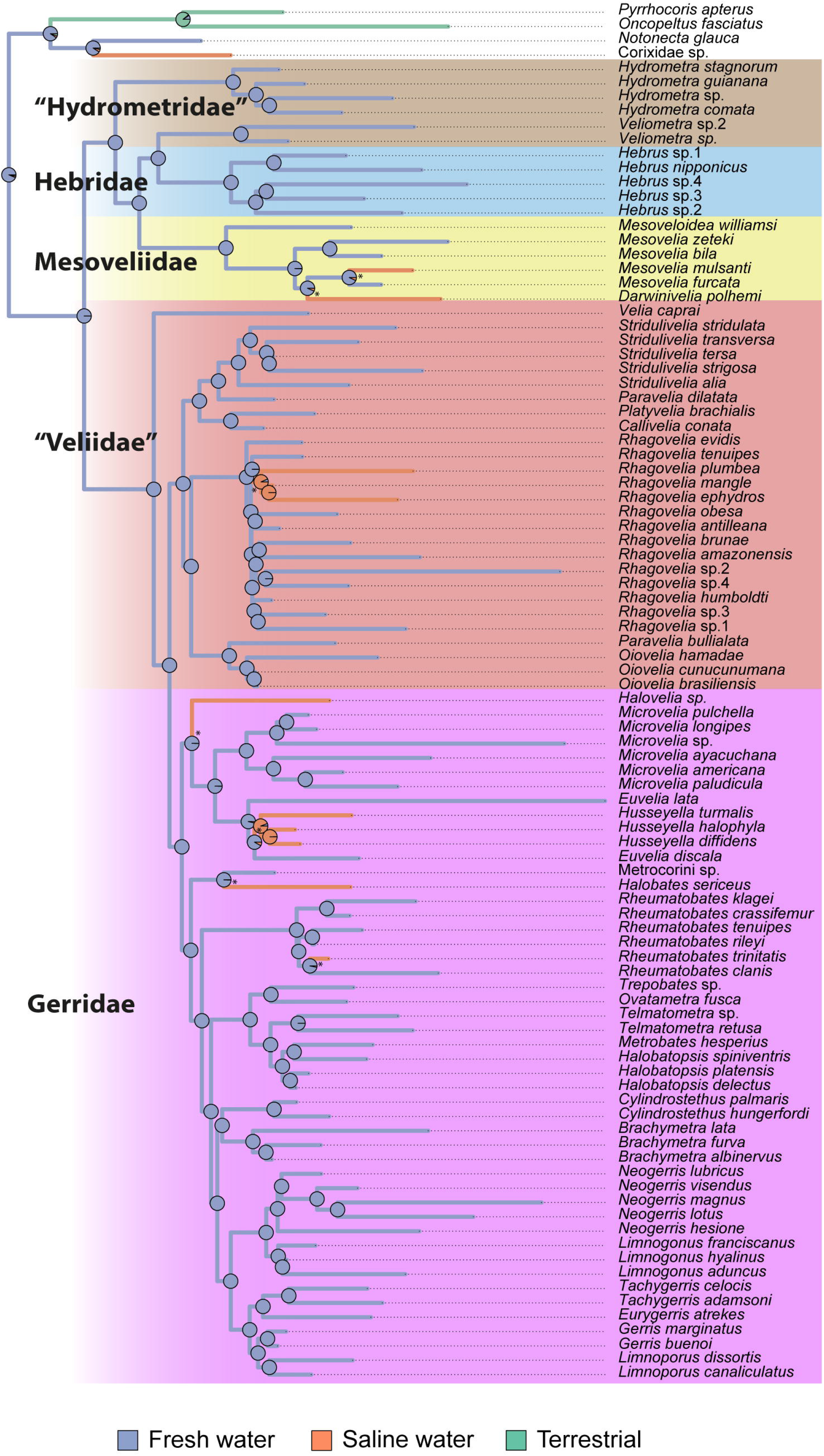
Ancestral character state reconstruction of habitat preference with regard to salinity. Pies represent the probability of ancestral state. Independent transitions marked with an asterisk (*).

### Evolution of wing polymorphism

In Gerromorpha, wing polymorphism is widespread and a variety of wing morphs, from complete absence (apterous) to long wing morphs (macropterous) have been described both within and across species (Figure 6). An intermediate morph (brachypterous) has been described in some species, but for simplicity this morph was not included in the reconstruction as amongst our sample it only occurs in few species also with both apterous and macropterous morphs. Among the species we sampled, 16 have exclusively long wings, 19 exclusively no wings, and the remaining species are polymorphic. *Cylindrostethus hungerfordi* was herein considered as entirely apterous, because only one macropterous specimen was recorded in the literature (Nieser 1970), which pends verification. The complete loss of wings is characteristic to marine species, which are represented in our sampling by the genera *Darwinivelia, Halovelia* and *Halobates* (Andersen 1982; Polhemus and Polhemus 2008). Among the 91 Gerromorpha species included in our phylogeny for which we had reliable wing morphs data, we reconstructed 7 independent events of wing loss, either at the genus level such as in *Veliometra, Darwinivelia, Halovelia, Halobates* and *Euvelia/Husseyella*, or at the species level such as in *Rhagovelia, Rheumatobates* and *Cylindrostethus*, where four, two and one species lost macropterous morphs respectively (Figure 6). Wing polymorphism, i. e. the presence of more than one wing morph in a single species, evolved twice independently within Mesoveliidae and in the common ancestor of (“Veliidae” + Gerridae) (Figure 6). Within the latter clade, the complete loss of wings evolved seven times and regains of a unique macropterous wing morph at least five times (Figure 6). These data suggest that the evolution of wing polymorphism is a dynamic process potentially influenced by the large heterogeneity of habitats, in terms of stability and seasonality, and the wide geographical distribution that these insects can occupy (Spence 1989; Harada and Numata 1993; Spence and Anderson 1994; Harada and Spence 2000; Fairbairn and King 2009).

**Figure 6:**
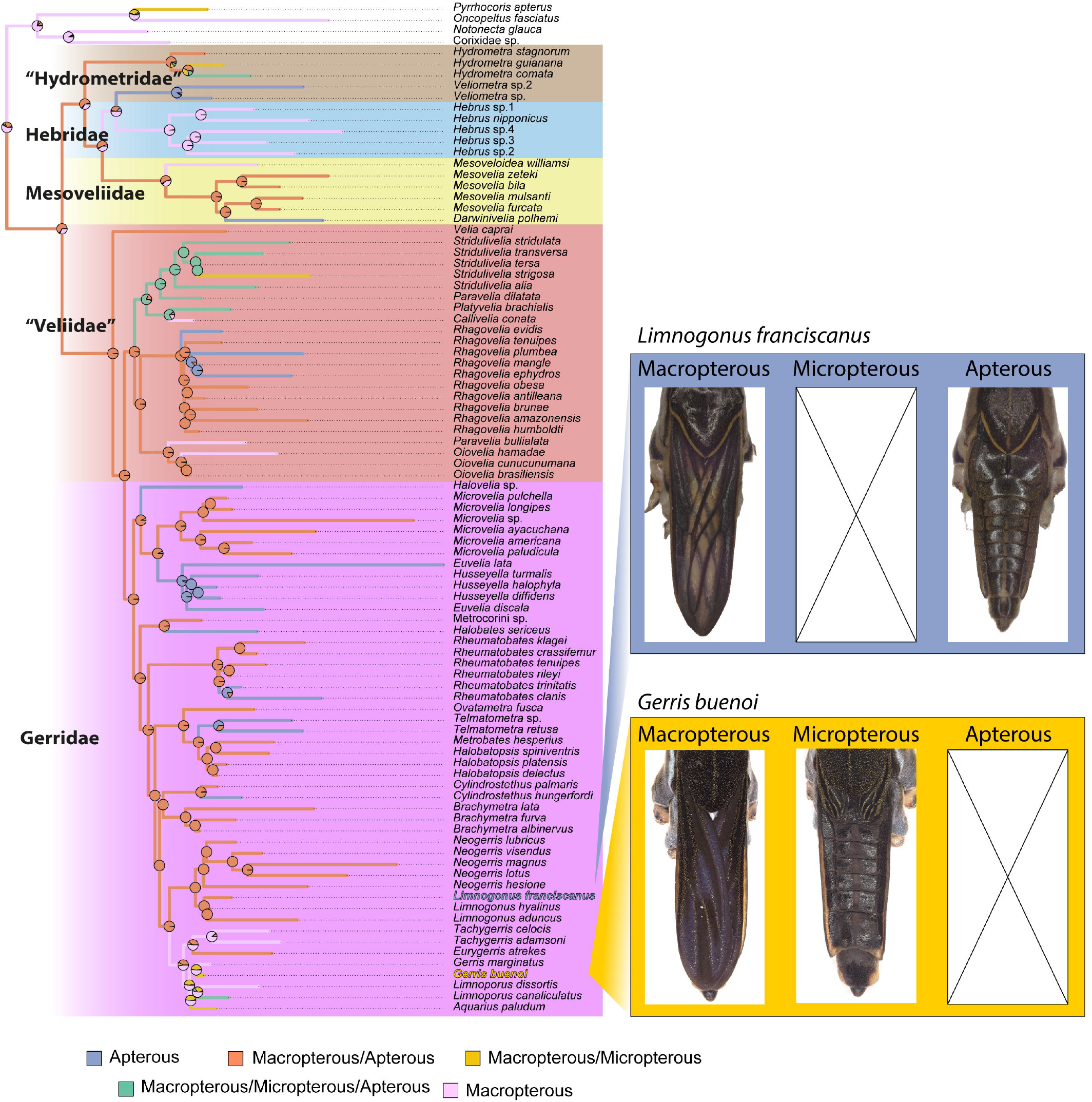
Ancestral character state reconstruction of wing polymorphism. Pies represent the probability of ancestral state. Images represent examples of wing polymorphism in two species: macropterous and apterous in *Limnogous franciscanus*, and macropterous and micropterous in *Gerris buenoi*.

## Discussion

Using about nine hundred transcripts as markers, we generated a new phylogeny of the Gerromorpha that resolved inconsistencies and uncovered strong support for an early split between the clade leading to the “Veliidae” + Gerridae and that leading to the “Hydrometridae” + Hebridae + Mesoveliidae. This change has important implications for analyzing and interpreting the evolutionary history of the group. We found that the evolutionary history of the Gerromorpha is rich in terms of transitions and expansions into new adaptive zones, enabled by important phenotypic changes to their appendages in association with new modes of locomotion on water. The multiple subsequent transitions to new adaptive zones may have been contingent on the initial event of water surface invasion common to all Gerromorpha.

### Evolutionary history of Gerromorpha

The results from our divergence time analysis suggest that the origin of Gerromorpha dates from the early Triassic, about ~20 MYA older than previous estimations using combined morphological and ribosomal data (Damgaard 2008a; Damgaard 2008b), but similar to other recent phylogenomic analysis (Johnson, et al. 2018; Wang, et al. 2021). Besides this divergence time difference, our phylogenomic approach using a large number of genetic markers, instead of classical ribosomal DNA phylogenies combined with morphological traits (Damgaard 2008a; Damgaard 2008b), provided three major outcomes. First, we could clearly define “Hydrometridae” as a basal clade from which (Mesoveliidae + Hebridae) split. This result contrasts with earlier analysis that placed Mesoveliidae as the earliest branching clade within Gerromorpha and sister to all other families of the infraorder (Damgaard 2008a; Damgaard 2008b), but is consistent with the latest reconstruction (Wang, et al. 2021). Second, we found that (“Veliidae” + Gerridae) diverged early in the evolution of Gerromorpha and is sister to (“Hydrometridae” + (Mesoveliidae + Hebridae)) rather than a later branching clade (Damgaard 2008a; Damgaard 2008b). These two results are inconsistent with the widely accepted hypothesis and our own previous results (Crumière, et al. 2016; Santos, et al. 2017; Vargas-Lowman, et al. 2019). However they are not unexpected, because: i) fossil records suggest that “Hydrometridae” and “Veliidae” probably appeared at a similar age, while Hebridae records are much more recent (Supplementary Material Table 3); ii) our increased number of species in the families “Hydrometridae”, Mesoveliidae and Hebridae allowed us to avoid long branch attraction artifacts that plagued our previous phylogenetic reconstructions; and iii) phylogenetic analyses focused on other heteropterans already suggested a similar evolutionary scenario for Gerromorpha (Johnson, et al. 2018; Wang, et al. 2021). Finally, our phylogenetic reconstruction supports the hypothesis that Microveliinae and Haloveliinae are subfamilies of Gerridae instead of “Veliidae”, which also holds up based on their tarsal segmentation and the morphology of the female gynatrial complex (Andersen 1982).

### Diversification of the legs in the Gerromorpha

The unique event of transition to water surface locomotion was accompanied by multiple phenotypic changes associated with exploiting surface tension, such as increased density of hydrophobic bristles (Andersen 1976), and generating efficient thrust on water (Hu and Bush 2010). We previously hypothesized that locomotion on water favors increased speed, which requires changes in locomotion parameters such as increased leg length and stroke frequency (Crumière, et al. 2016). This change was brought about through a gain of a new expression domain of the gene *Ubx* in the midlegs, in addition to its ancestral expression in the hindlegs (Khila, et al. 2009; Refki, et al. 2014). This role of Ubx had been possible after the initial gain of expression in the midlegs at the base of Gerromorpha, which enabled the increase in midleg length. The current ancestral character state reconstruction shows that leg length increased significantly at the base of the phylogeny, coinciding with the transition to the water surface (Supplementary Figures 2 to 7). This state of leg length was retained in, or regained by, lineages occupying the ancestral adaptive zone at the margin of water bodies (Figure 3). The “Veliidae” and Gerridae split early during the diversification of the infraorder and the split coincided with the evolution of rowing, enabling access to yet another adaptive zone composed of open waters (Figure 3). It appears that this new zone had many open niches as these two families now account for about 80% of all gerromorphans (Polhemus and Polhemus 2008). The change to the rowing body plan evolved at the base of these two taxa (Figure 3) and brought about deep changes in their locomotion behavior (Andersen 1982; Crumière, et al. 2016). Rowing gerromorphans generate fast and efficient thrust with significantly low frequency of strokes compared to walking Gerromorpha (Crumière, et al. 2016). This change is thought to allow these animals to sustain fast and efficient movement and expand into the vast surfaces of open water zones ranging from puddles to oceans (Cheng 1976; Andersen 1999).

Subsequent to the acquisition of rowing, certain taxa further evolved evolutionary innovations that allowed the transition to numerous other new niches. The family “Veliidae” contains about 60 genera and some 900 species, and the genus *Rhagovelia* alone accounts for almost half of the species count of the entire family (over 400) (Andersen 1982; Polhemus and Polhemus 2008). *Rhagovelia* spp. use rowing and all members of the genus possess a plumy fan in each midleg that allows generating efficient locomotion on fast flowing streams (Andersen 1982). The plumy fans are exclusive to the genus *Rhagovelia* and their evolution is likely to have enhanced their ability to survive in these fast-flowing streams. Four lineages (*Oioivelia, Paravelia, Callivelia* and *Microvelia*) made the move back from the rowing to the walking body plan, and also back to the margins of water bodies (Figure 3) (Andersen 1976; Crumière, et al. 2016). Interestingly, among these lineages, *Oiovelia* became specialized in foamy habitats that form on the margins of streams, typically trapped by fallen tree trunks or other obstacles (Rodrigues, et al. 2014). Therefore, phenotypic changes, and the associated genotypic changes, not only can allow lineages to acquire large new adaptive zones, but access to these also opens the possibility for further transition in a succession of niches where each new transition depends upon the previous.

In our character state reconstruction, it is noticeable that the modifications of male hindlegs into grasping traits evolved multiple times independently but exclusively in the lineages that diverged after the acquisition of rowing as a mode of locomotion (Figure 4). It is possible that the evolution of rowing, which almost exclusively relies on the midlegs to generate movement (Andersen 1976; Hu, et al. 2003; Crumière, et al. 2016), may have reduced constraints on the shape of the hindlegs due to biomechanical and hydrodynamics requirements no longer imposed by the ancestral walking mode. This change of functional emphasis on the role of the hindlegs may have freed these structures to evolve under sexual selection pressures. Coincident with the move to this two-dimensional habitat are high intersexual interactions that may increase sexual conflict, which in turn favors the evolution of males grasping traits such as modified hindlegs (Rowe, et al. 1994; Arnqvist 1997).

It is possible that the initial genetic changes involving Ubx, which shaped the legs in association with water surface locomotion, have provided the developmental genetic context for further modification of the legs under sexual conflict again in association with Ubx function. The finding in four tested lineages, out of seven, that the Hox gene *Ultrabithorax* is always required for male hindleg modification indicates that few developmental genetic paths are available for sexual conflict to drive male modifications. However, while the modifications are highly similar in species that possess spines (*e. g. Rhagovelia* and *Microvelia*), they are divergent from others (*Rheumatobates* in particular) where the elaborations involve deeper changes in the shape of the segments and the presence of sets of large setae along the proximo-distal axis. This is an indication that the developmental genetic pathways controlled by Ubx might diverge among these lineages, resulting in phenotypic differences of the modifications of male legs.

### Specialization on saline habitats and dispersal through flight

In our reconstruction, the ancestral habitat of the Gerromorpha is fresh water, consistent with previous reconstructions (Cheng 1976; Andersen 1999). Within Gerromorpha, the transition to saline water evolved throughout the phylogeny in at least seven independent instances out of the ten we sampled (Figure 5). Lineages transiting from freshwater to salty habitats faced a significant environmental change that is expected to affect fluid balance and concentration of electrolytes, which in turn is expected to pose significant challenges to osmoregulation. Common garden experiments testing tolerance to change in salinity revealed that many marine and fresh water lineages die quickly, with spasms and other signs of disrupted osmoregulation, when exposed to fresh and salty waters, respectively (Kishi, et al. 2006). The lineages that have made this transition must have undergone, independently, a series of physiological changes to regulate body fluid homeostasis. The nature of osmotic adaptation to brackish and saline water and the genetic mechanisms underlying these adaptations remain unknown in Gerromorpha.

Wings in Gerromorpha are important for dispersal through flight. Dispersal however is heavily influenced by environmental factors such as day/night cycles, habitat integrity or population density. In species with multiple wing morphs, population composition will depend on a combination of both genetic factors and the environment (Järvinen and Vepsäläinen 1976; Roff 1986; Simpson, et al. 2011), such as population density (Harada, et al. 1997; Harada and Spence 2000), food scarcity (Harada and Nishimoto 2007), salinity (Kishi, et al. 2007), temperature (Harada, et al. 2011), dryness (Harada 1998), photoperiod (Harada and Numata 1993), and habitat state (Cunha, et al. 2020). Wing development is an energy consuming process that is known to tradeoff with fertility (Hyun and Han 2021) and the presence of two or more wing morphs is widespread in Gerromorpha. Apterous, micropterous and brachypterous individuals can be considered as a single functional category with the inability to disperse by means of flight. Our phylogenetic reconstruction revealed that the presence of multiple wing morphs is ancestral in Gerromorpha, with highly dynamic patterns of gains and losses as the lineage diversified within the various aquatic habitats (Figure 6). Interestingly, the complete loss of wings, and therefore the ability to disperse, is tightly associated with the transition to salty water bodies, as there is an over-representation of wing loss in halophilic species. The loss of wings has been linked to the stability of marine water environments, which may have favored investment in reproduction rather than dispersal. Nevertheless, some fresh water lineages, such as *Veliometra* and *Euvelia*, are also exclusively wingless, raising the question as to what characteristics of their habitat or reproductive behavior may have driven wing loss.

## Conclusions

Next-generation sequencing technologies, by decreasing costs and increasing the amount of information generated has been key to our ability to reconstruct the phylogenetic history of previously understudied lineages. The use of transcriptome datasets is even more affordable and realistic as whole genome sequencing of a large number of species is still costly and requires accurate genome annotation. Drawbacks of phylotranscriptomics based on next generation sequencing include difficulties to obtain complete gene sequences, limited data for weakly expressed or non-expressed genes and uncertainties in gene number mainly due to the presence of multiple isoforms. However, many of these potential problems can be mitigated with a proper design of sampling, good preservation of specimen, enough sequencing depth, and third generation sequencing technologies. Transcriptome-based phylogenomics have been already used with success in the study of Hemiptera (Johnson, et al. 2018), true water bugs (Wang, et al. 2021) and, as presented here, in the semi-aquatic bugs. This approach has been critical in accurately rebuilding the phylogeny and allowed a reliable mapping and reconstruction of the evolutionary history of traits associated with their diversification. The data presented here constitute a valuable resource to the community interested in the study of phenotypic evolution.

## Methods

### Samples collection and culture

Natural populations were collected mainly during fieldwork in French Guyana and Brazil (Supplementary Material Table 1). Additional samples were obtained from short trips to Canada and USA or from personal communication. Field samples were classified and stored in individual tubes per species in RNA*later*™ Stabilization Solution (Thermo Fisher Scientific). When possible, lab populations were generated from this initial natural population. The bugs were isolated by species and maintained in a bug room at 25□°C and 50% humidity in water tanks and fed on crickets. RNA was extracted either from lab populations or from natural population samples if lab breeding was unsuccessful.

### Transcriptome sequencing

Poly-A RNA was isolated from embryos, nymphs and/or adults of ninety-two Gerromorpha, two Notonectidae and one Pyrrhocoridae (Supplementary Material Table 1). Total RNA was extracted using TRIzol (Invitrogen) according to manufacturer’s protocol. A first batch of libraries were sequenced using Hiseq 2500 paired end 100 nucleotides while a second batch was sequenced using Hiseq X teen paired end 150 nucleotides (Supplementary Material Table 1).

### Transcriptome assembly

Transcriptomic reads were assembled using Trinity 2.5.1 (Grabherr, et al. 2011) and long orfs with minimum protein length of 100 aa were extracted using TransDecoder 3.0.1 (Hass and Papanicolau). We used cd-hit (Li and Godzik 2006; Fu, et al. 2012) with a sequence identity threshold of 0.995 and a word size of 5 to reduce the number of redundant transcripts.

### Other transcriptomes

In addition to our 95 newly sequenced species, we included in our analyses the transcriptomes of six additional Gerromorpha as well as three close species available. *Limnoporus dissortis* (Gerromorpha) transcriptome was sequenced using 454 technology (Armisén, et al. 2015). *Gerris buenoi* (Gerromorpha) transcriptome was obtained from its annotated gene set (Armisén, et al. 2018). The transcriptomes of *Gigantometra gigas* (Gerromorpha), *Hebrus nipponicus* (Gerromorpha) and *Stenopirates* sp. (Encicocephalomorpha) were seguenced independently using Illumina Hiseq 2000 paired end 90 nucleotides, while *Halovelia* sp. (Gerromorpha) was sequenced using Illumina Hiseq 2000 paired end 150 nucleotides (Wang, et al. 2016). Finally, *Oncopeltus fasciatus* (Pentatomomorpha) gene transcripts were obtained from official gene set v1.1 (Panfilio, et al. 2019). *Hoplitocoris* sp. (Enicocephalomorpha) and *Ceratocombus* sp. (Dipsocoromorpha) transcriptomes were downloaded from (Johnson, et al. 2018).

### Contaminants removal

A list of sequence identifiers (gi) for bacteria, other, unclassified and viruses, where e-fetched from NCBI database to detect potential contaminants. Nucleotide sequences were then extracted from ‘nt’ database. We also included all *Gryllus* and *Acheta* gi sequences to this database of potential contaminants. We did so because brown crickets are used as a food source for all Gerromorpha species bred in the laboratory. A total of 75,358 transcripts amongst all the species were hit (threshold < 1e-5), classified as contaminants and removed from further analysis.

### Orthologous gene clusters

Transcripts from coding proteins were aligned all-against-all using BLASTP (Altschul, et al. 1990; Altschul, et al. 1997) masking low complexity regions and threshold of 1e-5. Output results were then clustered for homologous sequences using the software package SiLiX 1.2.11 which implements an ultra-efficient algorithm based on single transitive links with alignment coverage constraints (Miele, et al. 2011). We ran SiLiX using stringent values of minimum 70% of identity and 80% overlap to define the clusters.

The 3,299,629 protein coding transcripts from all species yielded a total of 3,079,991 clusters, including singletons. Clusters where then filtered to remove those that fail to contain at least 80% of the species. A total of 3,169 clusters were retained. In the event of multiple sequences present for a given species in the cluster, we applied a two-step selection process using a custom PERL script. First, we selected for each species the transcript with cumulative best scores against the transcripts of other species in a all-against-all BLASTP. Manual check of the results showed the existence of some chimeric clusters due to long conserved domains. To resolve this, in a second step we sorted the transcripts by value, selected the transcripts with highest and lowest cumulative score to start and assigned the rest of transcripts to one or the other based on their hit p-values. This effectively divided the transcripts in two lists. We selected the longest list and repeated the same process again. Finally for each missing species in the list we selected the best hit transcript in the cluster. This method allowed us to define the more conserved transcripts for each species and manual check showed that it solved the problems of chimeric clusters.

Manual check of clusters alignments also showed the presence of gaps that might be filled with other transcripts of the cluster. This situation might be due either a real gene split in two, or most probably, due to a *de novo* transcriptome assembly artifact. In any case, to fix this situation we developed a custom PERL script which looked for alignment gaps longer than 50 amino acids in each species. Then it looked for other transcripts of the same species in the cluster which hits the gap region with a maximum e-value of 1e-20. A maximum overlap of 20 nucleotides between both transcripts was allowed. When a suitable filling transcript was found we joined the nucleotide sequences of both transcripts.

Final result consists of a list of 3,169 clusters that are present in at least 80% of the 104 species.

### Phylogeny reconstruction

Prior to phylogeny reconstruction each of the 3,169 transcript clusters were aligned using MAFFT (Katoh and Standley 2013) and we used GBLOCKS (Castresana 2000) to retrieve conserved codon regions with half gap position allowed. We concatenated the blocks with size > 50% and performed SRH test on each cluster to verify if our data is stationary, reversible and homogeneous (Naser-Khdour, et al. 2019). A total of 894 gene clusters pass the SRH test at position 1 and 2. To reconstruct a phylogeny we ran IQTREE (Minh, et al. 2013; Nguyen, et al. 2015) using codon position 1 and 2 accounting for 415,432 nucleotides (out of 623,148 nucleotides in 894 gene clusters), independent automatic model selection (best-fit models per partition listed in Supplementary Material Table 4) and 1,000 ultrafast bootstrap trees to provide robustness estimates for the resulting maximum likelihood tree.

### Phylogeny reconstruction using BUSCOs

To validate our orthologs clustering protocol and posterior phylogeny reconstruction we used BUSCOs results to build clusters of single copy genes. A total of 1,870 clusters containing sequences for at least 80% of the species were recovered (out of 2,510 BUSCOs). Those clusters were then used as input in our PERL scripts to first, select the best transcript for duplicated BUSCOs, and second to use the gap filling script to improve fragmented BUSCOs alignments. After aligning and retrieve conserved codon regions, a total of 524 gene clusters passed the SRH test at position 1 and 2 accounting for 203,750 nucleotides. Finally, phylogeny reconstruction using IQTREE resulted in a tree consistent with our initial approach using a larger number of markers, and which support all our findings (Supplementary Figure 8).

### Fossil calibration

Estimated age of four fossils of extinct Gerromorpha species, two extinct Nepomorpha species and two extinct Heteroptera were used to calibrate divergence time at various points of our phylogenetic tree (Supplementary Material 3). To date the phylogeny, we use MCMC implementation in BEAST2 (Bouckaert, et al. 2019) using accompanying GUI BEAUTi to build the launch file. To increase the site coverage, we retained only positions present in 98% taxa which accounted for 74 823 nucleotides. We used previous IQTREE phylogenetic tree as starting tree and ran 200,000,000 generations using a Birth Death Model sampling every 1,000 generations. A final tree with median ages was constructed using Treeannotator (part of BEAST2 package) with a 10% burnIn and analyzed with Figtree (Rambaut) (Supplementary Figure 9). Root age was estimated at 277.4 Mya (95% CI 254.53-304.12 Mya). We replicated the analysis in a new run with consistent results (Supplementary Figure 10).

### Ancestral character state reconstruction

Reconstruction of ancestral character states was performed in Rstudio using ape and phytools package (Revell 2012) to do stochastic character mapping (Huelsenbeck, et al. 2003; Paradis, et al. 2004) for 1,000 simulations using “ER” model.

### Nymphal RNAi

Double-stranded RNA (dsRNA) was produced for *Ultrabithorax* gene (*Ubx*) as described in (Refki, et al. 2014). T7 PCR fragments were amplified from complementary DNA (cDNA) template using forward and reverse primers both containing the T7 RNA polymerase promoter. The amplified fragments were purified using the QIAquick PCR purification kit (Qiagen, France) and used as a template to synthesize the dsRNA as described in (Refki, et al. 2014). The synthetized dsRNA was then purified using a RNeasy purification kit (Qiagen) and eluted in Spradling injection buffer (Rubin and Spradling 1982) in a 2 to 3 μg/μl concentration. For primer information, see (Refki, et al. 2014). Nymphal injections were performed in parallel in nymphs with either *Ubx* dsRNA or injection buffer as negative controls. Nymphs were reared in water tanks and fed with crickets until they developed into adults. We then compared adults obtained from *Ubx* dsRNA injection to those obtained from injection of buffer injection alone.

## Supplementary material

Scripts and proposed pipeline can be found here: https://gitlab.com/davidarmisen/transcriptomic-based-phylogeny-in-gerromorpha

Aligned sequences of 894 gene clusters used for phylogeny reconstruction can be found in the same site.

## Declarations

### Ethics approval and consent to participate

Not applicable

### Consent for publication

Not applicable

### Availability of data and material

BioProject ID: PRJNA774202

### Collection permits

SISBIO permits 47975-3 and 47975-4.

## Acknowledgements

We thank Lois Taulelle and Hervé Gilquin for providing access to computing resources in the Pôle Scientifique de Modélisation Numérique (PSMN) at the ENS Lyon; and Jakob Damgaard for the discussions on Gerromorpha phylogeny and evolution over the years. We thank François Bonneton for discussion and comments. We also thank Qiang Xie for data sharing and discussion.

## Supplemental Figure legends

**Supplementary Figure 1:** Phylogeny reconstruction with IQTREE using our final list of 894 clusters of orthologous genes for a total of 623,148 aligned nucleotides across the 104 species. Support values are indicated in each node.

**Supplementary Figure 2:** Ancestral character state reconstruction of foreleg to body length ratio in males.

**Supplementary Figure 3:** Ancestral character state reconstruction of midlegs to body length ratio in males.

**Supplementary Figure 4:** Ancestral character state reconstruction of rear leg to body length ratio in males.

**Supplementary Figure 5:** Ancestral character state reconstruction of foreleg to body length ratio in females.

**Supplementary Figure 6:** Ancestral character state reconstruction of midlegs to body length ratio in females.

**Supplementary Figure 7:** Ancestral character state reconstruction of rear leg to body length ratio in females.

**Supplementary Figure 8:** Phylogeny reconstruction with IQTREE using 524 gene clusters build from BUSCOs results.

**Supplementary Figure 9:** Dated phylogenetic tree. Nodes contain divergence dates and purple bar represent 95% confidence interval.

**Supplementary Figure 10:** Replicate dated phylogenetic tree. Nodes contain divergence dates and purple bar represent 95% confidence interval.

## Supplemental Table legends

**Supplementary Table 1:** Resume table with information about the new 95 species sequenced. It includes information about the isolate (natural population or lab population), collection date, geographical location, GPS coordinates, developmental stage sequenced, sex and sequencing technology.

**Supplementary Table 2:** Resume table with the number of sequenced reads, assembled transcripts and genes for each of the 95 newly sequenced species. Additionally, we also include BUSCOs stats for all the 104 species transcriptomes used, regardless of the source.

**Supplementary Table 3:** Fossil records used to set the calibration points to date the divergence time in the phylogenetic tree of Gerromorpha.

**Supplementary Table 4:** Best individual model for each charsets using BIC.

