## Supplementary figures and images for "Transcriptome-based phylogeny of the semi-aquatic bugs (Hemiptera: Heteroptera: Gerromorpha) reveals patterns of lineage expansion in a series of new adaptive zones"

### Supplementary Figure 1

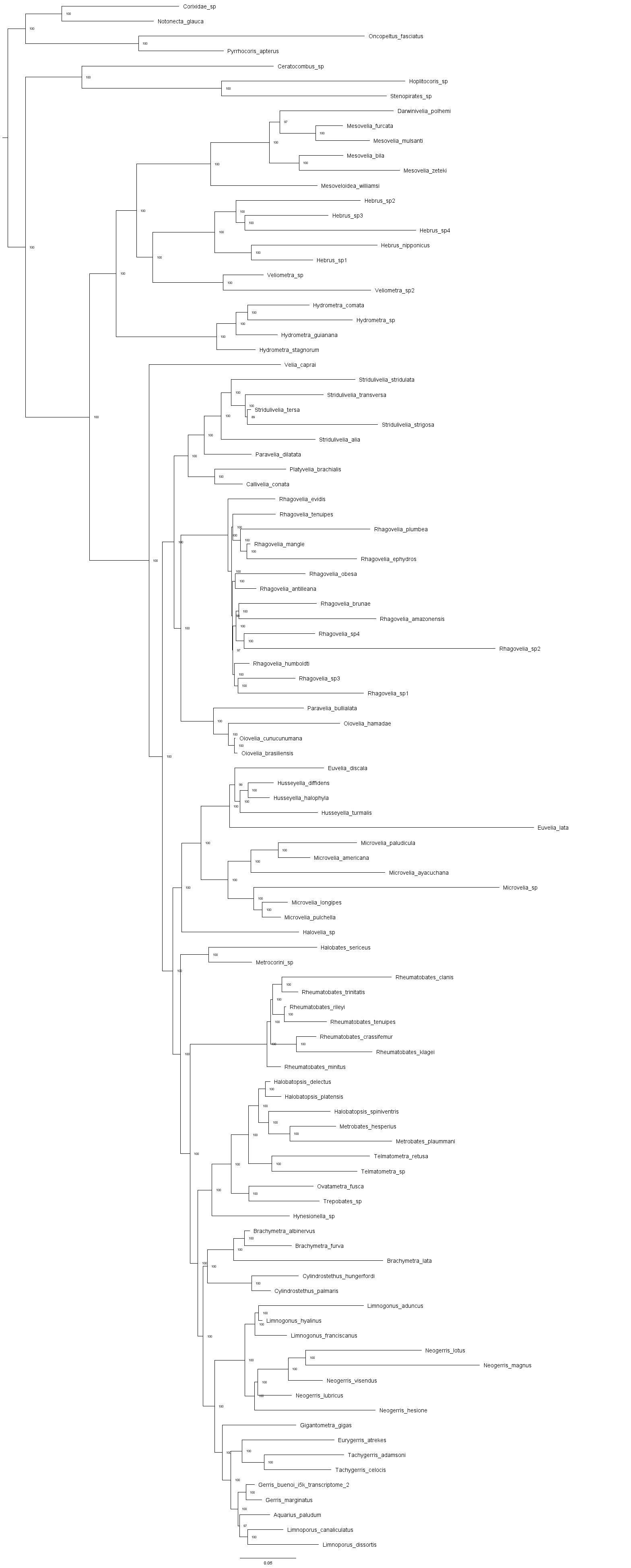

### Supplementary Figure 2

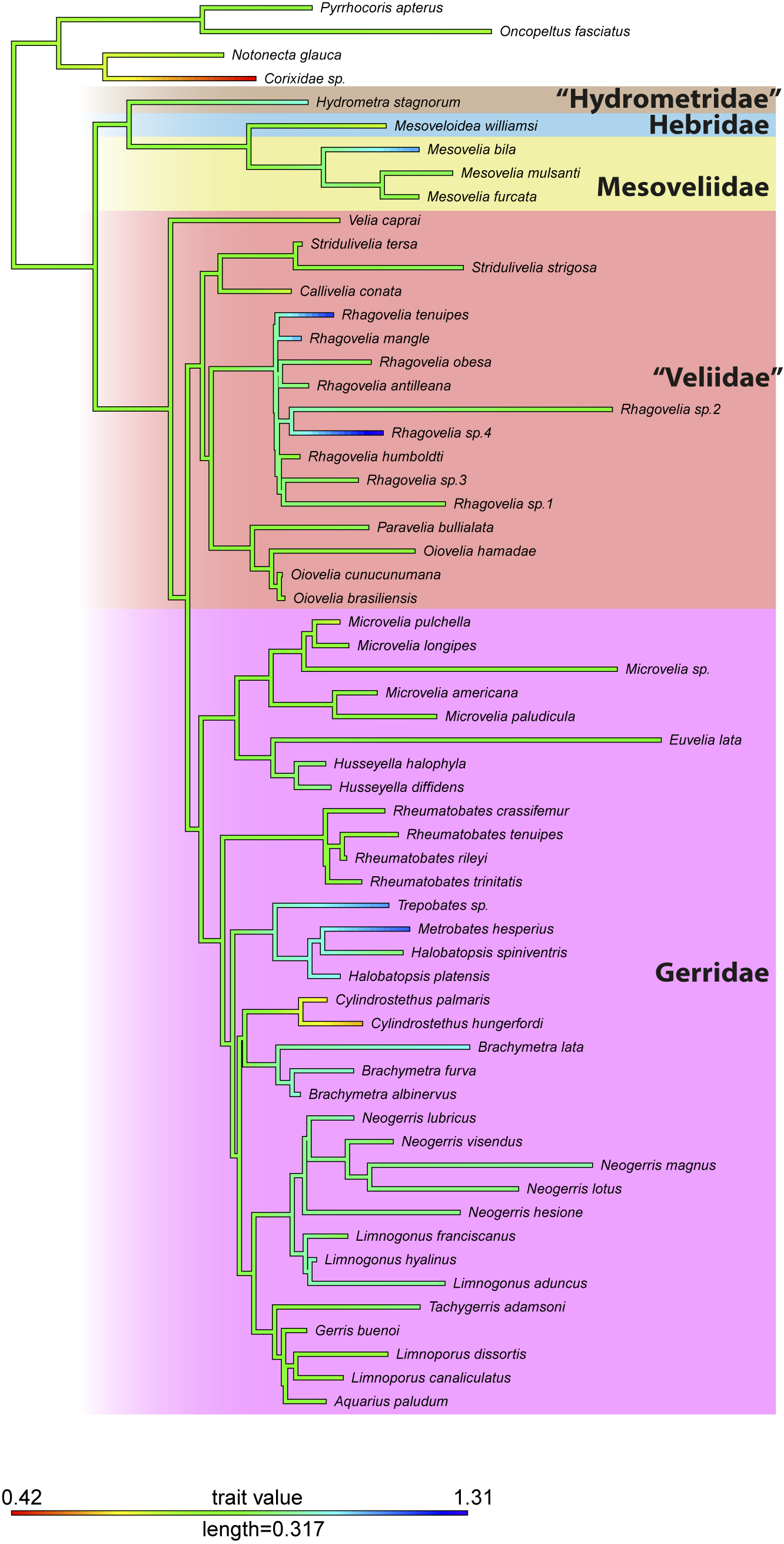

### Supplementary Figure 3

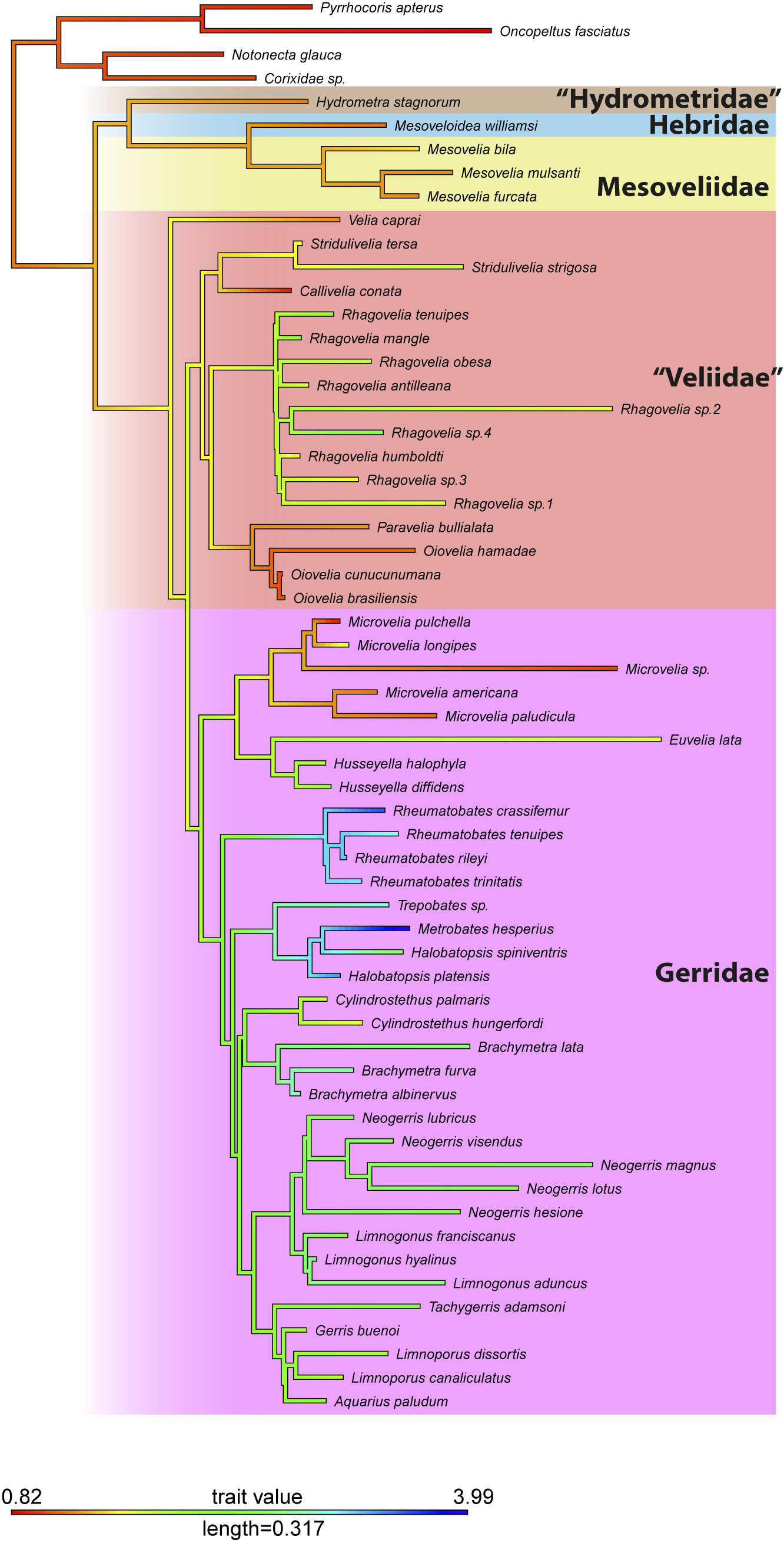

### Supplementary Figure 4

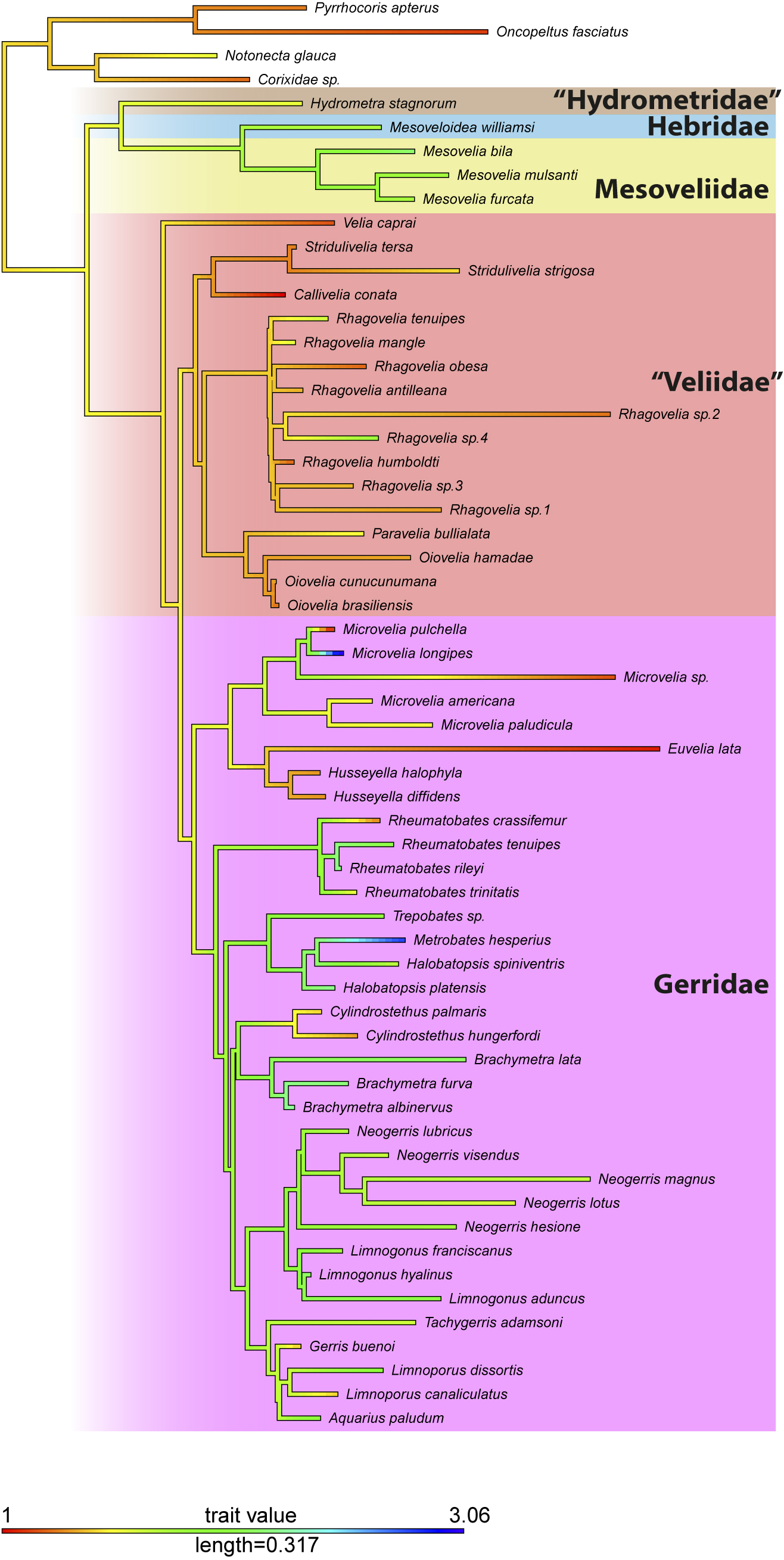

### Supplementary Figure 5

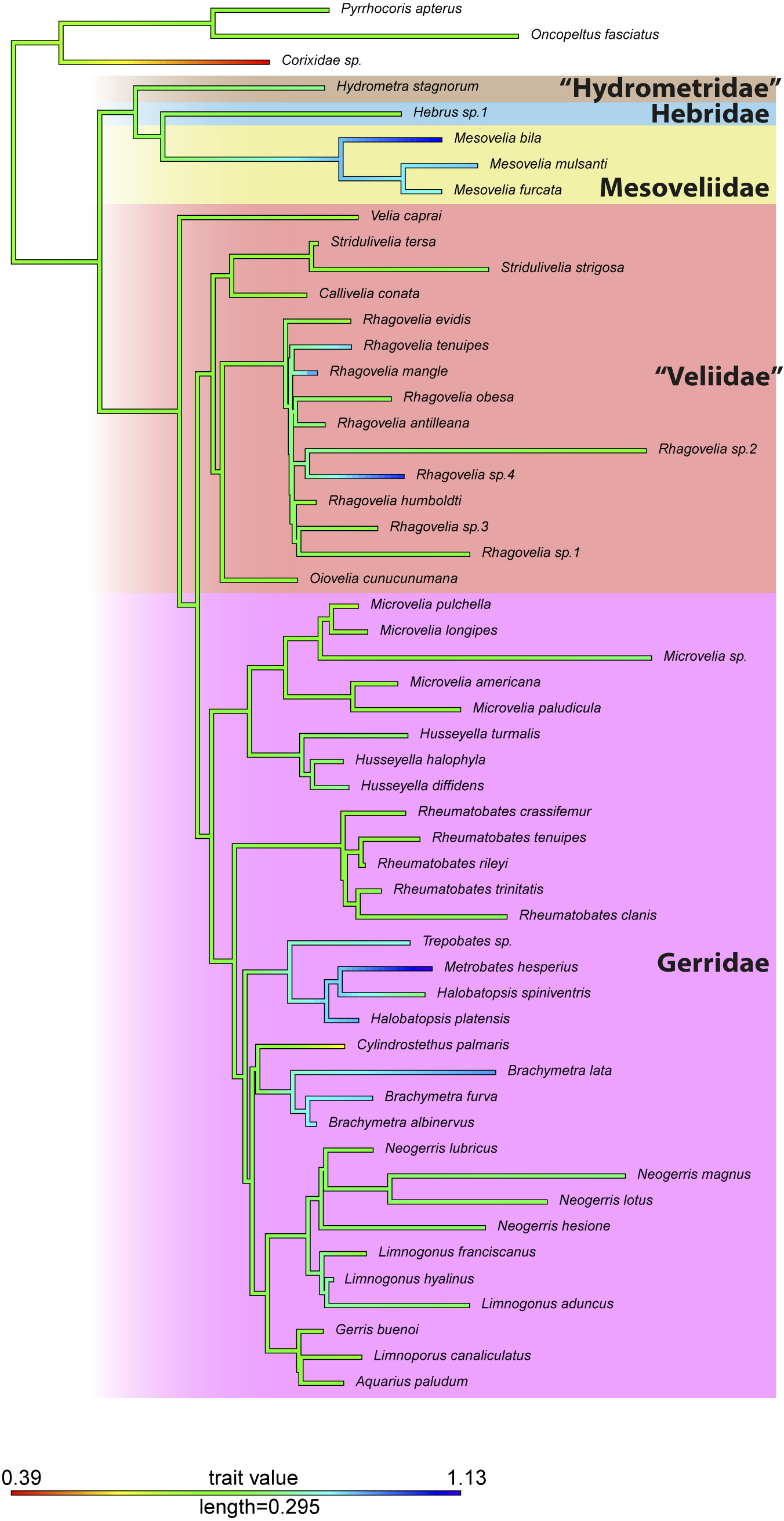

### Supplementary Figure 6

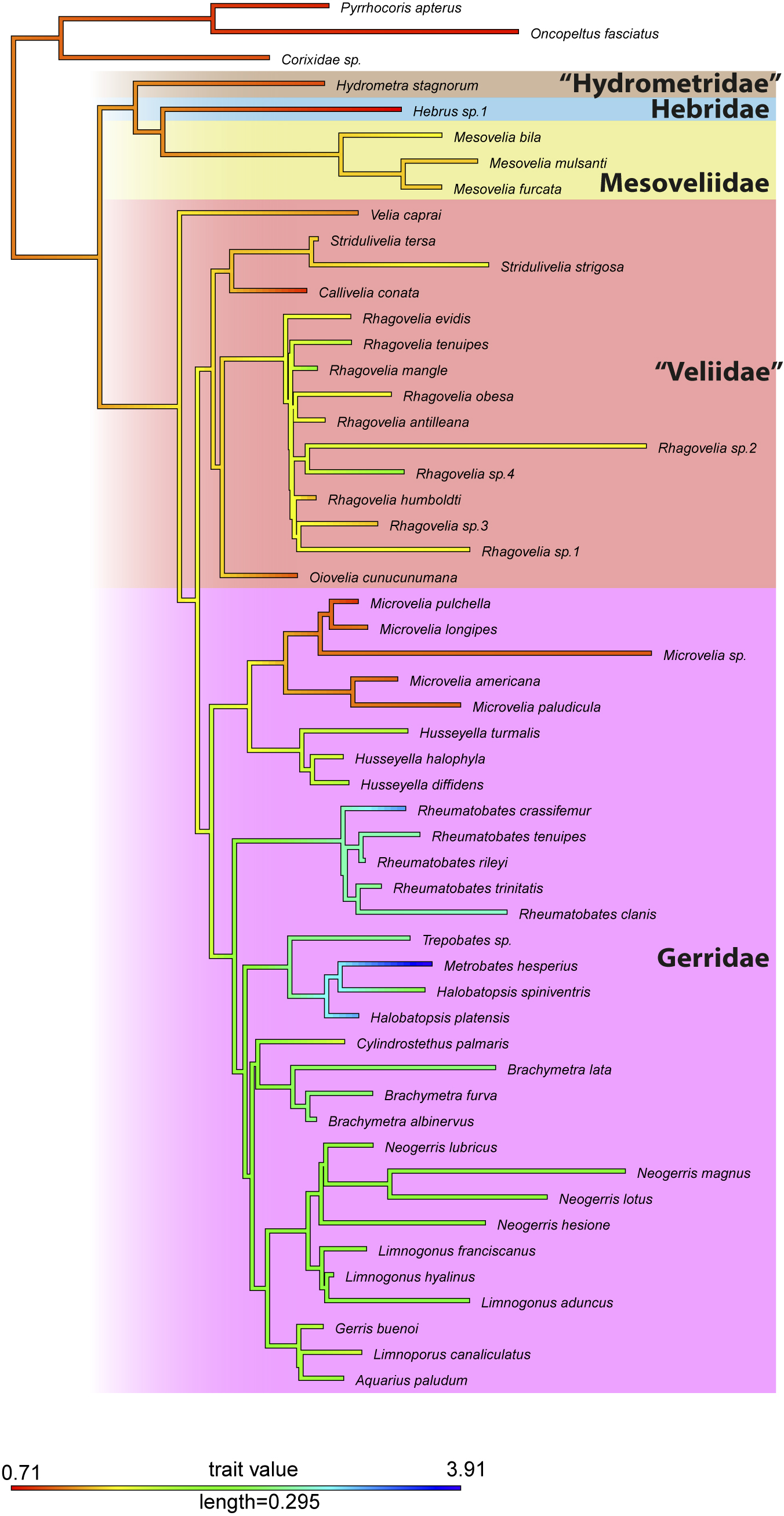

### Supplementary Figure 7

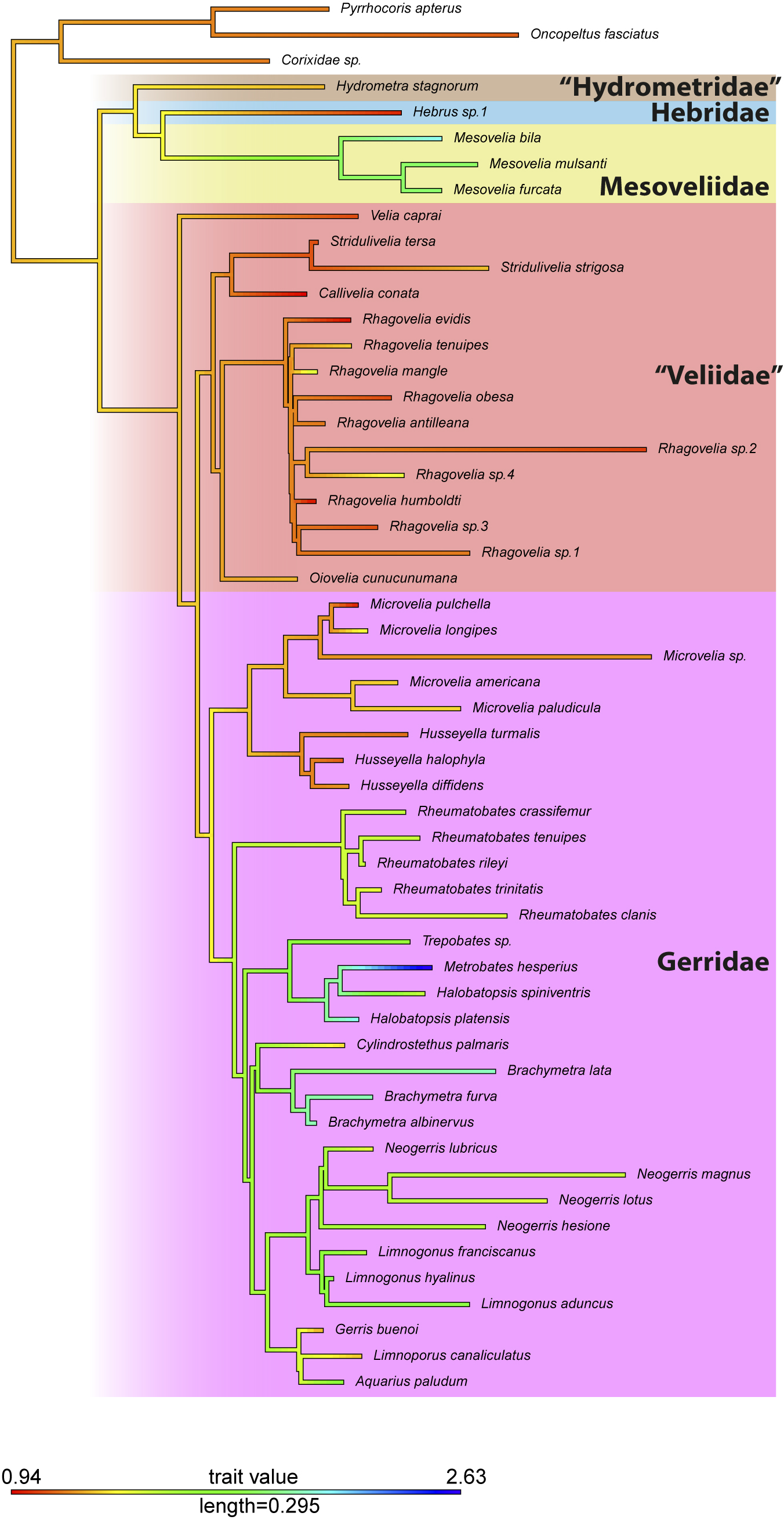

### Supplementary Figure 8

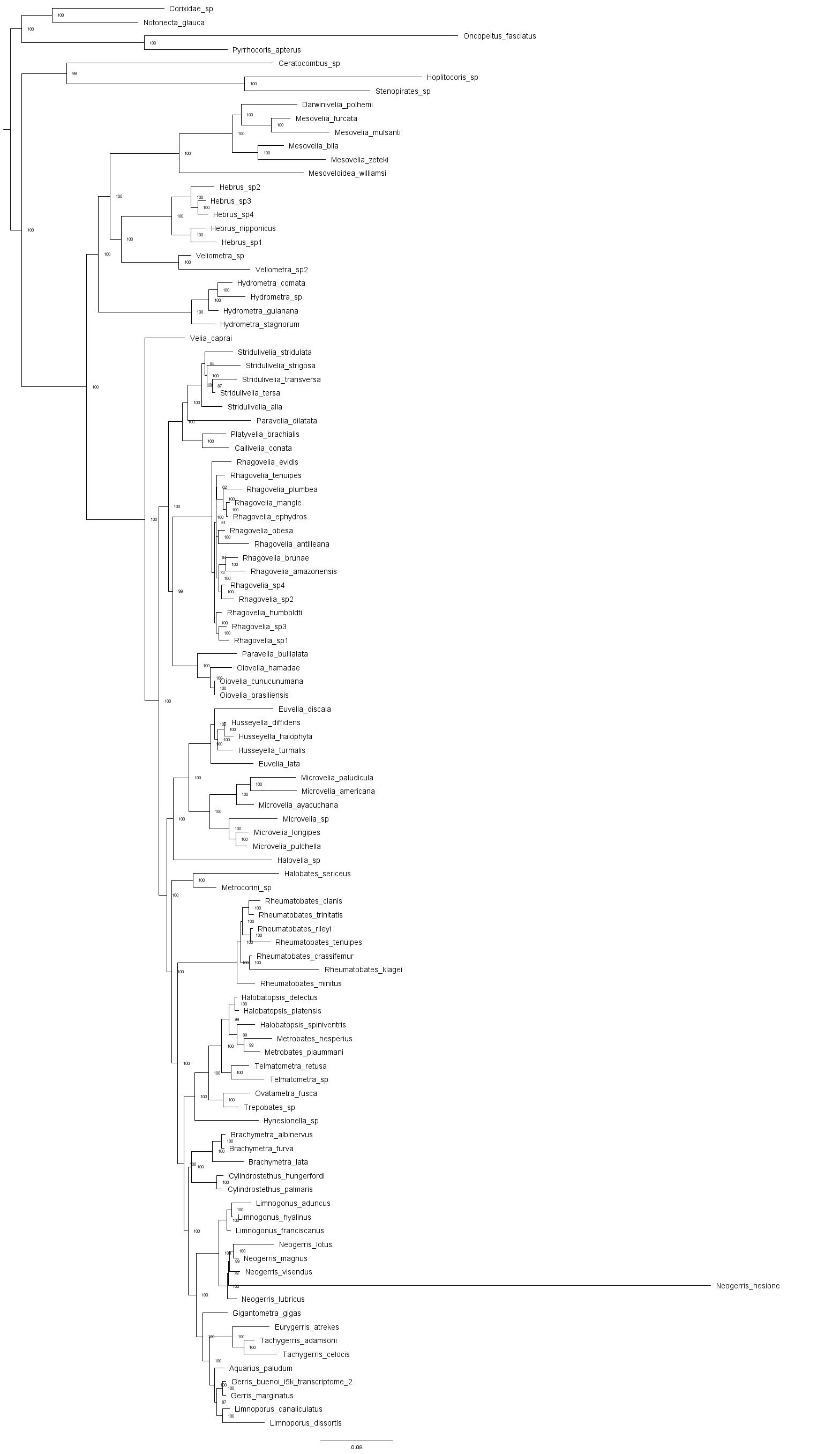

### Supplementary Figure 9

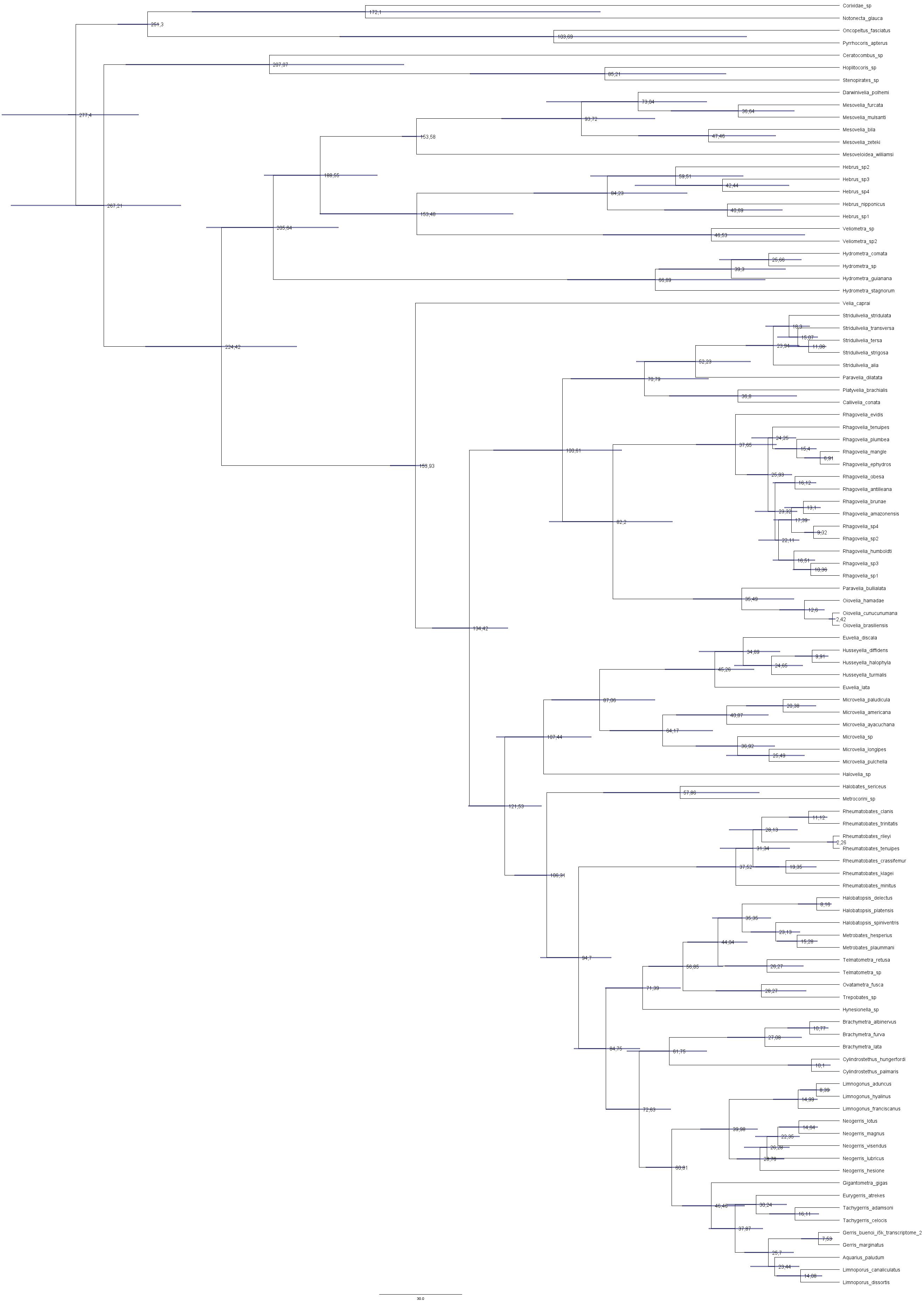

### Supplementary Figure 10

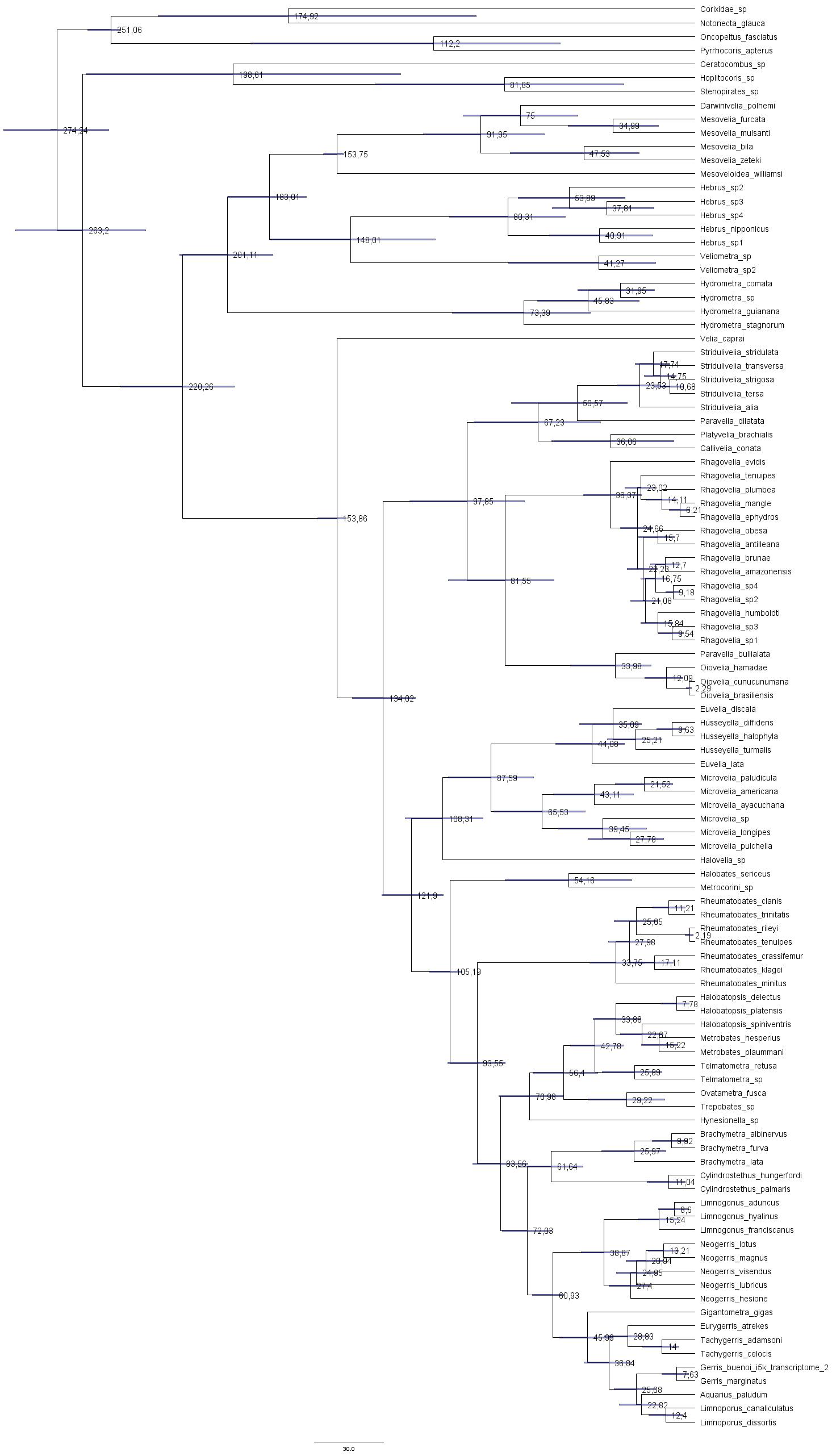
